# What’s left from the brew? Investigating residual barley proteins in spent grains for downstream valorization opportunities

**DOI:** 10.1101/2025.03.26.645420

**Authors:** Simon Gregersen Echers, Rasmus K. Mikkelsen, Naim Abdul-Khalek, Lucas Sales Queiroz, Timothy J. Hobley, Benjamin Schulz, Michael Toft Overgaard, Charlotte Jacobsen, Betül Yesiltas

## Abstract

Brewer’s spent grain (BSG) is the major side-stream from beer production but remains highly underutilized. While the direct use of BSG as a food ingredient is limited due to subpar techno- functionality, the vast amounts and fairly high protein content of up to 30% makes it a high potential source for production of protein-based ingredient by valorization through e.g. enzymatic hydrolysis. However, little attention has been put towards the protein-level composition of BSG, which is essential for developing hydrolysis strategies for improving functionality in a targeted manner. Here, we present an in-depth characterization of the BSG proteome and investigate dynamic proteome changes from malting and mashing in the initial phases of beer production. We show dynamic and selective changes in the proteome across the different process steps, where 29% of reproducibly identified proteins display differential abundance. BSG represents a significantly higher proportion of intracellular protein compared to both barley and malt and has a nutritionally favorable amino acid composition. The major constituent of the BSG proteome is B3-Hordein, constituting more than 30% of the BSG protein. Moreover, we find that a large proportion (> 45%) of the BSG protein is associated with potential food safety concerns, being classified as potential allergens and antinutritional factors. Our analysis emphasizes the need for downstream processing of BSG to produce safe and functional food ingredients, while also providing protein-level insights for development of targeted hydrolysis strategies to achieve this.

**Highlights:**

- An optimized sample preparation for proteomics analysis has been developed
- 29% of barley proteins are differentially abundant across malting and mashing
- B3-Hordein is the major protein in BSG with an abundance over 30%
- BSG contains a high content of potential allergenic and antinutritional proteins
- A protein-level basis for targeted downstream processing of BSG is presented

## 1. Introduction

Brewer’s spent grain (BSG) is the major side-stream from beer production, representing up to 85% of the total waste from the brewing industry [1]. With an annual production of around 200 billion liters of beer from the global brewing industry, around 40 million tons of BSG is generated [1,2]. While BSG has applicability in production of e.g. energy, bioethanol, and paper, it is still considered an underutilized resource [3]. Currently, most BSG is used in cattle feed or end up in landfills [4,5]. To facilitate a better utilization and increase the value of BSG, the desire for its application in food products has increased within the last decade and several companies actively use BSG as an ingredient in products such as bakery items and snacks [6]. However, BSG inclusion levels are typically below 20% due to a lack of compatible techno-functional and sensory properties, resulting in a negative impact on e.g. texture and color, ultimately leading to consumer rejection [6,7]. One of the main reasons is the high content of residual fiber (30-50% w/w) [7,8] whereof up to 50% is constituted by indigestible fibers such as arabinoxylan [9], limiting direct use. One strategy to improve functionality and thereby applicability of BSG is to enrich protein and reduce fiber content through downstream processes [4,5]. In this way, the typical protein content of 19-30% (w/w) in BSG [7,8] can be enriched up to 85% [10] and almost full protein extractability is achievable [11,12]. However, achieving both simultaneously is not readily achievable and usually requires rather harsh treatments, including acids, organic solvents, and elevated temperatures, which further impair the techno-functionality, particularly solubility, of the BSG protein [1,5,13].

One way to valorize BSG and improve its techno-functionality is through enzymatic hydrolysis. In fact, enzymatic treatment has previously been shown to increase protein yield from BSG compared to e.g. alkaline extraction [14,15], but also to improve functional and bioactive properties of BSG proteins [5,16,17]. Previously, potent antioxidant peptides from barley glutelins (i.e. hordein) have been identified [18]. Moreover, combining enzymatic hydrolysis with emerging green extraction methods has also been shown to improve both yields and functionality [19]. Recent advances in prediction of peptide-level properties, such as antioxidant activity [20] and emulsification [21,22] has facilitated a fundamentally different approach in hydrolysate production, governed by mass spectrometry (MS)-based proteomics combined with bioinformatic data analysis and prediction of peptide functionality using artificial intelligence [23]. This has led to discovery of many novel functional and bioactive peptides [24–27] and has also illustrated the potential for valorization of several sustainable protein sources [28–30]. The concept was recently demonstrated in potatoes [31], yielding hydrolysates with improved surface-active properties *in vitro* that were furthermore able to e.g. improve stability of perishable fish oil through encapsulation [32]. But to adapt such a strategy, a detailed characterization of the BSG proteome is required.

BSG is the side-stream from the initial phase of beer brewing prior to fermentation [33]. Barley seeds are allowed to partially germinate during malting, which upregulates a range of hydrolases facilitating the degradation of both proteins and carbohydrates within the seed endosperm storage [34]. Following milling, malted barley is infused with hot water during mashing, where solubilized protein and starch is degraded to peptides and free amino acids (AAs) as well as fermentable sugars, respectively. Hops is added to the resulting liquid phase (wort) and used as the basis for the fermentation to produce beer, while the solid residue constitutes the BSG. While many barley proteins ultimately end up in the final beer product [33], a substantial amount of protein ends up in the underutilized BSG. Barley, as well as the proteome dynamics during malting and mashing, has previously been investigated to understand the underlying enzymatic machinery of the processes [35–39]. Such insight can enable optimization both in terms of process parameters but also in cultivar selection and development, where key enzymes can be enriched and/or optimized to facilitate a more efficient process [40–42]. But to date, most focus has been on the wort and final beer product [33]. Very limited effort has been made to describe the protein-level changes during mashing and how this determines the final protein composition in BSG. This knowledge gap hinders development of targeted processing strategies to valorize this massive side stream.

In this work, we present a deep characterization of the industrial barley “Planet”, its corresponding malted form, commonly used in Danish brewing, and use a conventional mashing procedure to produce representative BSG. These streams of the pre-fermentation brewing process were compared using a range of different methods to investigate bulk changes. Using bottom-up proteomics, we investigate the protein-level changes at each stage of the process, dive into the BSG proteome to identify the most abundant proteins, and use this insight to evaluate the potential benefits and challenges of using BSG as a food ingredient.

## 2. Materials and methods

### 2.1. Production of brewer’s spent grains

Brewer’s spent grains (BSG) was produced by conventional mashing, as previously described [43]. Briefly, 50 g of malted barley (*Hordeum vulgare*, var. Planet) (Viking malt, Vordingborg, Denmark) was milled using a 0.7 mm DLFU disc mill (Bühler, Skovlunde, Denmark). The grist was added to 150 mL 0.3 mM CaSO4 (in dH2O) pre-heated to 65 °C in a mashing bath (Locher Labor + Technik GmbH, Berching, Germany) and rested for 50 min. Subsequently, the temperature was increased at a rate of 1 ℃/min to 74 ℃ and rested for 10 min. After addition of 150 mL dH2O (pre-heated to 74°C), the mash was rested for an additional 10 min before being cooled to 25°C and subsequently, 100 dH2O was added for a final grist-to-water ratio of 1:8. The wort was removed by first running the mash through a common strainer and then through Whatman prepleated 320 mm filter paper (Cytiva, Marlborough, USA). The first filtered 100 mL was reused to wash the BSG and returned. All retained BSG was pooled and dried overnight at 60 °C.

### 2.2. Crude protein characterization

The two BSG batches, along with the malted barley, were analyzed to establish crude characteristics of the streams. Unprocessed barley was obtained from the same batch and variety as the malted barley used for BSG production (Viking malt, Vordingborg, Denmark).

#### 2.2.1. Dry matter

Dry matter was determined gravimetrically. 2 g of sample was weighted off in pre-weighted and dried beakers and placed in a heating cabinet at 105°C for 24 hours. The samples were cooled in a desiccator and weighted with 4 decimals. All samples were measured in triplicates.

#### 2.2.2. Crude protein content

The crude protein (CP) content was estimated based on nitrogen (N) content by DUMAS combustion using a Rapid Max N exceed (Elementar UK Ltd, Handforth, England). A nitrogen-to-protein conversion factor of 6.25 was used to calculate protein content. All determinations were performed in triplicate.

#### 2.2.3. Amino acid analysis

Amino acid composition was determined based on method described by Ghelichi *et al*. [44]. Briefly, approximately 30 mg of sample was hydrolyzed with 6 M HCL at 110°C for 18 h. Samples were cooled and filtered through 0.22 µm cellulose acetate syringe filters and subsequently pH adjusted and diluted by 200 mM potassium hydroxide and 100 mM ammonium acetate. Amino acid composition was analyzed with LC-MS (Agilent 1100 series, LC/MSD Trap) with a BioZen Glycan LC Column. Glutamine, asparagine, tryptophan and free cysteine cannot be determined using this method. Glutamine and asparagine are converted into glutamic acid and aspartic acid, respectively, and are reported as a combined sum. Disulfide-bound cysteine (cystine) is not degraded and is included in the analysis. Measurements were carried out in triplicates.

#### 2.2.4. SDS-PAGE

One-dimensional SDS-PAGE was performed under reducing conditions, as previously described [45]. Briefly, analysis was conducted using SurePAGE 4-20% polyacrylamide gels (Genscript, USA) and a Tris-MOPS SDS Running Buffer system (Genscript). For solid samples, aliquots of cryogenically milled grains were mixed with 4x SDS sample buffer (50 mM Tris pH 6.8, 2% SDS, 10% glycerol, 0.02% bromophenol blue, 12.4 mM EDTA, and 50 mM DTT) to a protein concentration of 4 mg/mL (based on CP) and subsequently incubated for 5 min at 95 °C. Afterwards, samples were allowed to cool, briefly centrifuged to sediment particles, and the supernatant recovered for analysis. For size estimation, 5µL PIERCE Unstained Protein MW Marker was loaded. Electrophoreses was carried out at 150 V for 40-50 min until the dye front reached the bottom of the gel. Gels were subsequently stained using Coomassie blue and imaged on a ChemDoc MP imaging System (BioRad, USA).

### 2.3. Proteomics analysis

To characterize the protein-level composition of the different biomasses, they were investigated using bottom-up proteomics (BUP). Prior to analysis, grain samples (barley and malt) were cryogenically milled using a mortar and pestle with liquid nitrogen. Milling was continued with addition of liquid nitrogen until a fine and homogenous flour was obtained. Cryo-milling was also performed on BSG to investigate any potential benefits in terms of protein extractability. Milled grains and BSG (milled and crude) were lyophilized to ensure a completely dry starting material prior to further processing.

#### 2.3.1. Protein extraction optimization

To optimize the sample preparation protocol in relation to protein extraction efficiency, the effect of combining focused ultrasound with the commercial iST for plant tissue kit (PreOmics, Germany) was initially investigated. For each sample, an amount corresponding to 800 µg protein based on CP was weighed off into Low-Bind tubes (Eppendorf, Germany) and 1 mL “Lyse” buffer was added. Tubes were vortexed and placed on a pre-heated Thermomixer (Eppendorf) and incubated at 95°C for 5 minutes at 1000 rpm. The solutions, and as much undissolved biomass as possible, was transferred to individual 1 mL AFA tubes (Covaris, USA) and the protein was extracted using a M220 focused ultrasonicator (Covaris). All samples were extracted through 0, 1, 2, 4, and 8 cycles of the protein extraction protocol specified by the manufacturer (peak incident power of 75 W, a duty factor of 10%, 200 cycles per burst, and 180 s per cycle at 6°C). After each of the indicated number of cycles, aliquots of the extracts were transferred to new Lo-Bind tubes and subjected to a secondary round of heat incubation on the thermomixer using the same conditions. Extraction efficiency was subsequently evaluated by SDS-PAGE analysis. To ensure comparability, SDS-PAGE analysis was performed using equivolumetric sample load of 30 µL, corresponding to 24 µg protein (based on CP analysis assuming full protein extraction). Samples were mixed with 4x sample buffer with 200 mM DTT in a 3:1 ratio, and subsequently prepared and analyzed as described above for solid samples.

#### 2.3.2. Sample preparation for mass spectrometry

Prior to analysis by mass spectrometry, protein was extracted based on the optimized parameters. For grain samples (barley and malt), two AFA extraction cycles were applied, while for both BSG samples, one cycle was performed. After extraction and additional heat incubation to ensure full protein denaturation, disulfide reduction, and thiol carbamidomethylation, extracted protein was digested and desalted, as described by the manufacturer. Briefly, 100 µL of the extract (corresponding to 80 µg) was mixed with 50 µL resuspended protease (Trypsin/LysC) and incubated for three hours on a thermomixer at 37°C and 500 rpm. Subsequently, the digested protein was transferred to iST cartridges and subjected to three rounds of washing (including “Wash 0” for plant tissue), two rounds of elution, and finally dried in a SpeedVac concentrator (Thermo-Fisher Scientific, USA). When dry, samples were resuspended in 70 µL “LC Load” and the peptide concentration was estimated by microvolume UV-spectroscopy (A280) using an SDS-11 FX (Denovix, USA) at standard settings (1 Abs = 1 mg/mL) and diluted to reach a final concentration around 0.2 µg/µL. Samples were stored at -18°C until analysis. All extractions were performed in triplicate.

#### 2.3.2. Bottom-up proteomics by LC-MS/MS

Shotgun BUP was performed on EASY nLC-1200 ultra-high-performance liquid chromatography system (Thermo) with ESI coupled to a Q Exactive HF tandem mass spectrometer (Thermo). For each sample replicate, 1 μg digest was loaded on a PEPMAP trap column (75 μm x 2 cm, C18, 3 μm, 100 Å) followed by reversed-phase separation using a PEPMAP analytical column (75 μm x 50 cm, C18, 2 μm, 100 Å). The mobile phase (solvent A (0.1 %(V/V) formic acid) and solvent B (80 %(V/V) acetonitrile, 0.1 %(V/V) formic acid)), was introduced through a stepwise gradient from 5 % to 100 % solvent B over 60 minutes. The analysis was performed using full MS/ddMS2 Top20 data-dependent acquisition with an MS1 scan range of 300-1600 m/z, positive polarity, and a default charge of 2. The MS1 resolution was set to 60,000, while the dd-MS2 resolution was set to 15,000. AGC target and maximum injection time were set to 1e5 and 45 ms, respectively. The isolation window was defined as 1.2 m/z, collisional energy as 28 eV, and the dynamic exclusion window as 20.0 s. Peptide match was set as “preferred” and “exclude isotopes” was enabled.

#### 2.3.3. Raw data processing

Raw LC-MS/MS data was processed with MaxQuant v.2.2.0.0 [46]. The data was searched against the Ensembl Plants [47] *Hordeum vulgare cv. Morex* IBSC_v2 database [48]. Methionine oxidation and protein N-terminal methylation were used as variable modifications while cysteine carbamidomethylation was included as a fixed modification. Data was analyzed using tryptic *in silico* digestion, allowing up to two missed cleavages. Minimum peptide size was defined as 7 amino acids and maximum peptide mass as 4600 Da. A false discovery rate of 1% was employed on both peptide- and protein-level. Match between runs and dependent peptides were enabled to boost identification rates.

The mass spectrometry proteomics data have been deposited into the ProteomeXchange Consortium via the PRIDE [49] partner repository with the dataset identifier PXD062029 and 10.6019/PXD062029.

#### 2.3.4. Downstream data analysis

Based on the incomplete protein annotation in the IBSC_v2 database, the nature of identified proteins was determined by BLAST analysis using Uniprot [50]. Briefly, the sequences of identified proteins were extracted from the IBSC_v2 protein fasta into a custom fasta file using a custom script in Python (v.3.8.8). Similarly, subproteome fasta files (e.g. differential or abundant proteins) were constructed for subsequent BLAST analysis. BLAST hits were evaluated by identity, and e-value with a preference given for proteins in a Uniprot barley proteome (UP001057469 or UP001177060). If no high identity and low e-value hits originated from a barley proteome, preference was given to proteins with a specifically annotated name (i.e. avoiding “Uncharacterized” and “Predicted” proteins) from closely related cereal species. For proteins annotated as “Uncharacterized” or “Predicted”, the protein with the highest homology and distinct annotation was used to indicate protein function.

##### 2.3.4.1. Inter-sample comparative analysis

For inter-sample comparison using label-free MaxLFQ [51] data, Mass Dynamics 2.0 [52] was employed. Missing values were imputed using the built-in “missing not at random (MNAR)” algorithm with a mean position factor of 1.8 and a standard deviation factor of 0.3. For pair-wise differential analysis, proteins were considered significantly and differentially abundant if the adjusted p-value was lower than 0.05 (i.e. false discovery rate (FDR) < 5%) and the fold change ratio was larger than two (i.e. log2(FC) > 1). Data was subsequently visualized in volcano plots using the threshold values for significance and fold change. For hierarchical sample and protein clustering in heatmap representation, a Euclidean distance of 4 and a row-wise Z-score normalization was applied. Only proteins identified as differential by the built-in ANOVA analysis were included in the heatmap representation.

##### 2.3.4.2. Intra-sample relative quantification

For intra-sample quantitative analysis, intensity-based absolute quantification (iBAQ) [53] was employed using the MaxQuant built-in option. Initially, all false positive and contaminant proteins were filtered. Next, proteins were filtered based on the requirement that a protein must be quantitatively identified in at least two replicates of at least one sample. To obtain the relative molar protein-level abundance, iBAQ intensities were normalized to the sum of iBAQ intensities within a given sample, as previously described [54]. To further segment identified proteins and focus on only the most abundant proteins, filters of 0.5% riBAQ and 1% riBAQ were applied. Statistical analysis of intra-sample quantification was performed as described below.

##### 2.3.4.3. Subcellular localization

Using the custom fasta with only identified proteins, subcellular localization was predicted using DeepLoc 2.0 (https://services.healthtech.dtu.dk/services/DeepLoc-2.0/) [55]. While DeepLoc potentially provides multiple probable subcellular origins, only the compartment with the highest probability was used to categorize proteins. Within each predicted compartment, the riBAQ of all associated proteins were summarized to obtain the grand distribution of proteins based on subcellular origin.

### 2.4. Statistical analysis

Data from crude protein characterization and intra-sample relative quantification by riBAQ (including subcellular localization) was analyzed to identify statistical differences using GraphPad Prism (v.10.0.2, build 232). Comparison of means from the different streams was performed using ordinary two-way ANOVA and Tukey’s multiple comparison testing, with a 95% confidence interval. Assumptions included Gaussian distribution of residuals, equal standard deviations between groups of unmatched replicates, and a single pooled variance. All p-values are reported as adjusted p-values to account for multiple comparisons.

## 3. Results and discussion

### 3.1 Characterization of protein content and amino acid composition

Based on initial crude characterization (Fig. 1), the barley and malted barley were overall found more comparable to each other than to BSG. The significantly lower DM content in barley (p < 0.0001) may reflect a higher water retention prior to malting (Fig. 1A, left). In contrasts, the crude protein (CP) increased significantly (p < 0.001) over each step of the BSG production process (Fig. 1A, right), almost doubling from barley (9.0% DM basis) to BSG (17.4% DM). This finding agrees with previous studies [7,56,57] and highlights the potential of BSG as protein-rich side-stream, which should be explored and exploited to a higher extent than it currently is.

**Figure 1:**
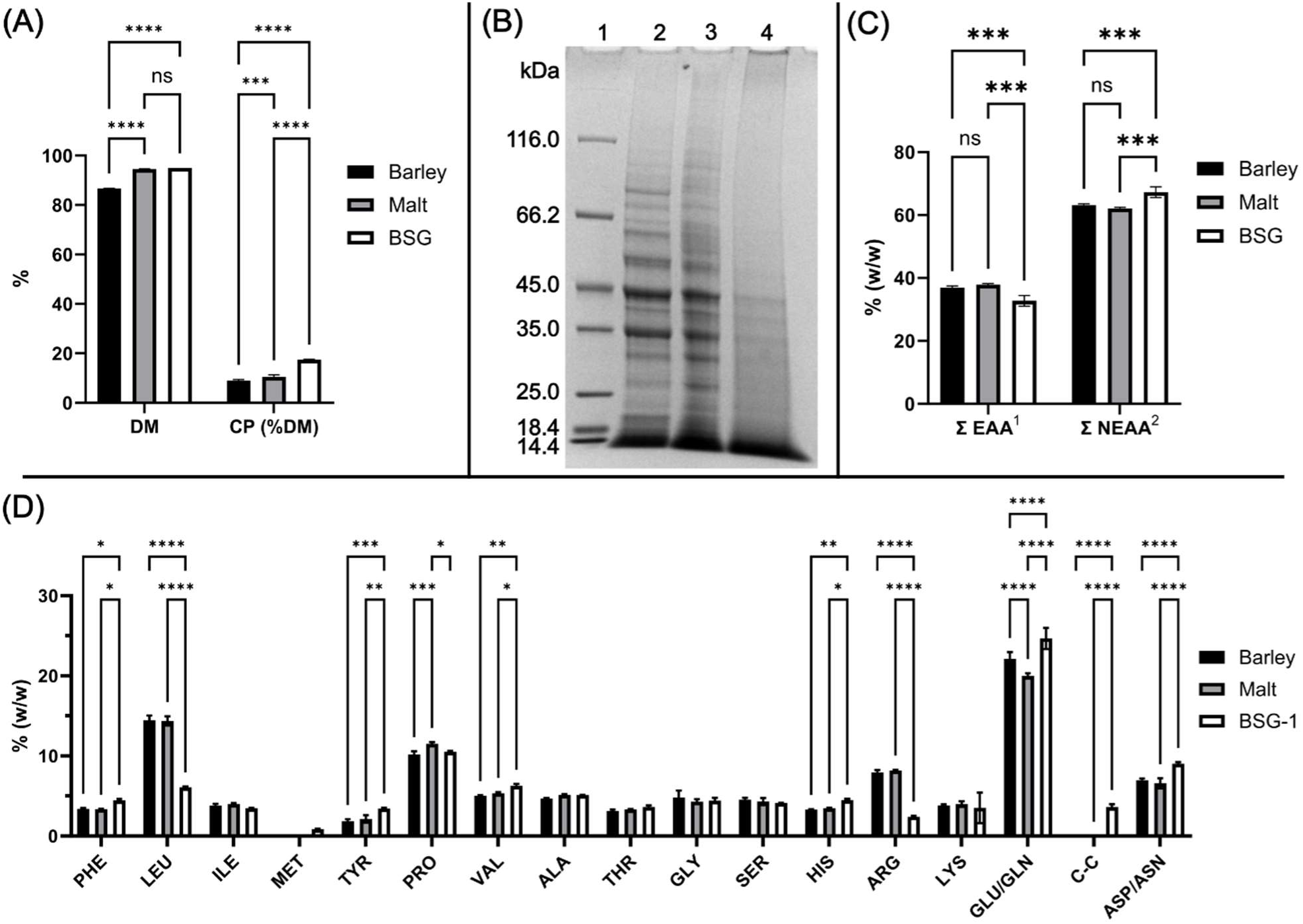
Initial characterization of the crude barley, malt and BSG. A) Dry matter (DM) and crude protein (CP) content. B) Reducing SDS-PAGE analysis of planet barley (2), malted planet barley (3), and the produced BSG (4). All samples were analyzed in crude form by directly solubilizing cryo- milled samples in SDS-sample buffer. As molecular weight marker, Pierce unstained marker was used (1) and mass (in kDa) is indicated. C) Sum of essential amino acids (Σ EAA) and non-essential amino acids (Σ NEAA) as per D) Amino acid analysis showing the distribution of individual amino acids as relative abundance (% w/w). Compositional analysis is indicated as means with the standard deviation (n = 3). Statistical analysis is performed as two-way ANOVA with significance level (from adjusted p-values) indicated by “ns” (p > 0.05), “*” (p ≤ 0.05), “**” (p ≤ 0.01), “***” (p ≤ 0.001), and “****” (p ≤ 0.0001). For the amino acid analysis (D), non-significant differences are not shown for simplicity. Numerical data can be found in Table S1.

The greater comparability between barley and malt, compared to BSG, is also reflected in both the protein profile by SDS-PAGE analysis (Fig. 1B) and the amino acid (AA) composition (Fig. 1C, D). Differences in visible protein bands and intensities are observed between barley and malt, particularly above 50 kDa, but the overall profile remains rather comparable. In contrast, BSG was void of most bands and only a few of the prominent bands (particularly in the range from 35 kDa to 45 kDa) were retained after mashing. In previous studies, prominent bands in this region have been ascribed to B- hordeins and embryo globulins [58,59]. Based on the amino acid profile, only the relative (% w/w) proportion of Pro and Glu/Gln differed significantly between barley and malt (Fig. 1D), and therefore the overall content of essential and non-essential amino acids (EAA and NEAA) showed no significant difference (Fig. 1C). In contrast, BSG showed a slight but significant (p < 0.001) decrease in EAA from 38-39% in barley and malt to 33% in BSG. While this represents a somewhat lower EAA content than e.g. milk protein (38%), it is still substantially higher than other plant-based proteins like oat (21%), wheat (22%), and soy (27%) and more comparable to egg (32%) and casein (34%), representing typical animal-based proteins [60]. Moreover, the EAA content was well above the 27% presented as the WHO/FAO/UNU amino acid requirements [61].

While the content of branched-chain AAs (BCAAs) Leu (6.1%) and Ile (3.5%) was lower in BSG than the comparable levels in barley and malt (approximately 14.4% and 3.9%, respectively), it still exceeds the WHO/FAO/UNU requirements of 5.9% and 3.0%, respectively. In contrast, the Val content was significantly enriched in BSG compared to barley and malt and a content of 6.3% is far beyond the required 3.9%. As such, BSG can be considered sufficient as a source of BCAAs. Based on the requirements, only Lys (3.5%) and Met (0.83%) were below the required levels, as BSG was also found rich in aromatic AAs. As Met is commonly considered in combination with Cys as sulphur-containing AAs, the sample cystine (C-C) content (3.6%) in BSG also makes it sufficient. As such, only Lys can be considered a limiting AA in terms of nutritional quality based on our analysis. Ultimately, these findings show that BSG may indeed be considered a promising source of nutritional protein for use in foods. They also emphasize that more effort and resources should be allocated towards finding better use of this by-product. Not only for nutritional purposes, but also for improving both economic and environmental sustainability of the brewing industry by converting waste into value.

### 3.2 Optimization of protein extraction for proteomic analysis

To gain deeper insights on the protein-level composition, we performed bottom-up proteomics (BUP) by LC-MS/MS. However, as the samples represent complex flours and because protein may still be retained in the recalcitrant plant matrix, thereby limiting accessibility, we initially explored protein extractability. While the employed iST kit is specifically developed for proteomics analysis of plant tissues [62], previous analyses have indicated that thermal and chemical treatment alone may not be sufficient to facilitate complete and representative protein extraction [63,64]. However, the combination of the kit with physical disruption methods, such as ultrasonication, has been shown to improve representative protein recovery [54]. Consequently, we attempted to combine the thermochemical lysis and extraction from the kit with adaptive focus ultrasound.

By subjecting all samples to multiple rounds of ultrasonication, we were able to evaluate extraction efficiency and protein quality by SDS-PAGE (Fig. 2). Application of other conventional quantitative protein assays (such as absorbance at 280 nm), and fluorochrome-based methods (such as Qubit), have previously been shown to be incompatible with the lysis buffer from the iST kit, due to either coextraction of non-proteogenic UV-active species or buffer incompatibility [54]. Consequently, SDS-PAGE analysis was applied for both qualitative and quantitative evaluation. From this analysis, it was evident that extraction from milled grain samples (barley and malt) benefitted greatly from the application of ultrasound (Fig. 2A). It was also evident that two extraction cycle appeared sufficient for both flours (Fig. 2A, lanes 4 and 9) and that excessive treatment with up to eight cycles (Fig. 2A lanes 6 and 11) had a detrimental effect for particularly higher MW proteins. BSG (Fig. 2B) appeared to not benefit from ultrasonication beyond one extraction cycle (Fig. 2B, lanes 3 and 9), but compared to no ultrasonication (Fig. 2B, lanes 2 and 8), a clear improvement was observed. Based on SDS- PAGE analysis, no apparent differences in the protein composition between crude and cryo-milled BSG were observed. The decreased requirement for mechanical force during extraction from BSG, compared to grain samples, is likely directly linked to the additional processing and breakdown of the recalcitrant matrix during mashing. Consequently, two cycles were employed for milled grain samples and one cycle for both BSG samples prior to BUP analysis. Compared to the SDS-PAGE analysis of the crude samples (Fig. 1B), the extracts displayed a high level of similarity in the protein profile, underlining the applicability of combining the iST kit and focused ultrasound for representative protein extraction.

**Figure 2:**
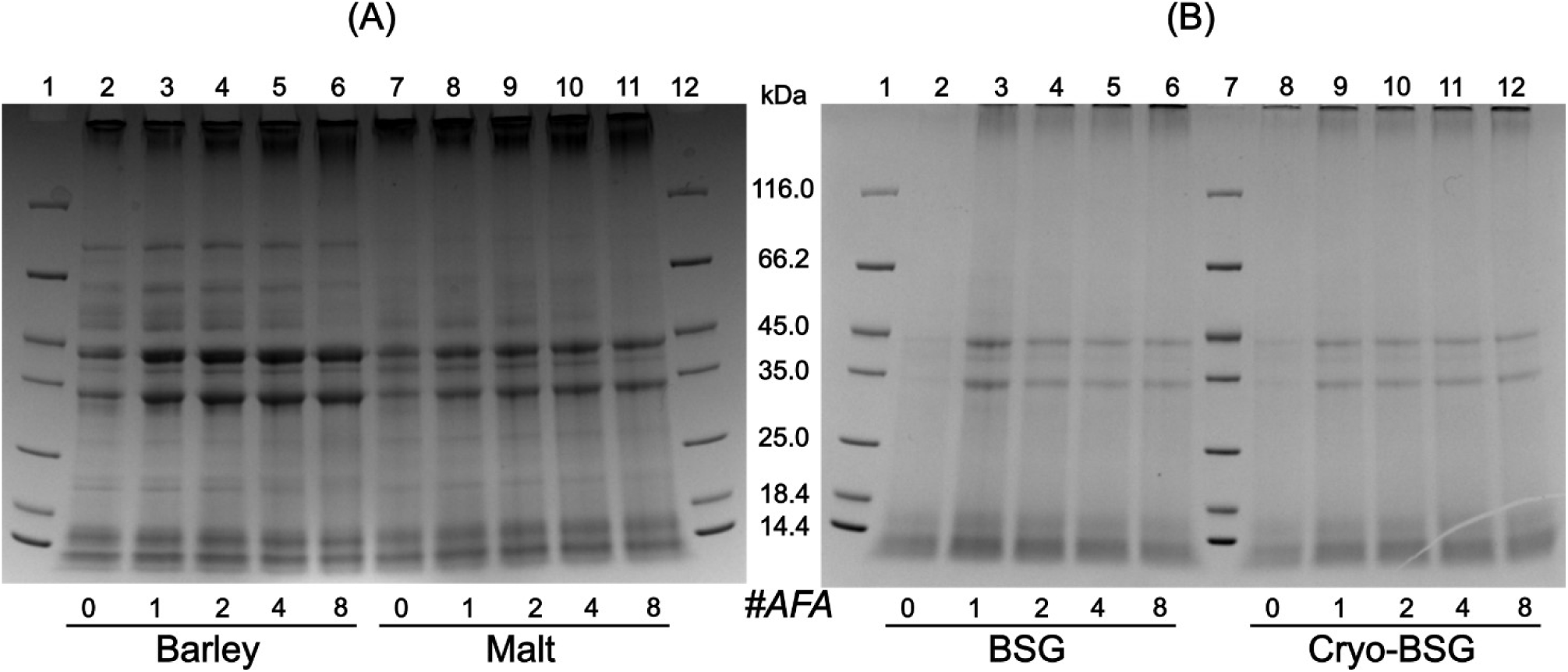
Evaluation of protein extractability by reducing SDS-PAGE analysis. Gels are shown for (A) barley (lane 2-6) and malt (lane 7-11) as well as for (B) BSG (lane 2-6) and Cryo-BSG (lane 8- 12). Protein was extracted using the iST plant tissue kit and in combination with 0, 1, 2, 4, and 8 AFA ultrasonication cycles (as indicated below each lane). As molecular weight marker, Pierce unstained marker was used (lanes A1, A12, B1, and B7)) and mass (in kDa) is indicated.

### 3.3 Overall protein-level changes during BSG production

Using the optimized protocols for sample preparation, we identified a total of 967 barley protein groups across all samples by LC-MS/MS-based BUP, following the removal of common contaminants and false positives. From these 967 protein IDs, 706 were quantitatively identified in at least two replicates from unprocessed barley, while 857 were found in the malted barley samples. In BSG, 647 protein groups were identified in at least two replicates from the crude sample, while 604 protein groups were reproducibly quantified in the cryo-milled sample. Across all samples, 945 protein groups (98%) were reproducibly found in at least one sample. Nearly half of the proteins (454) were identified in at least two replicates of all samples, while 660 proteins were shared between barley and malt (Fig. 3A). The proportion of proteins reproducibly identified in malt but not in barley (200) was substantially larger than the reverse (49). 196 proteins found in malt were not identified reproducibly in any BSG sample. These observations illustrate a dynamic process from barley over malt to BSG.

**Figure 3:**
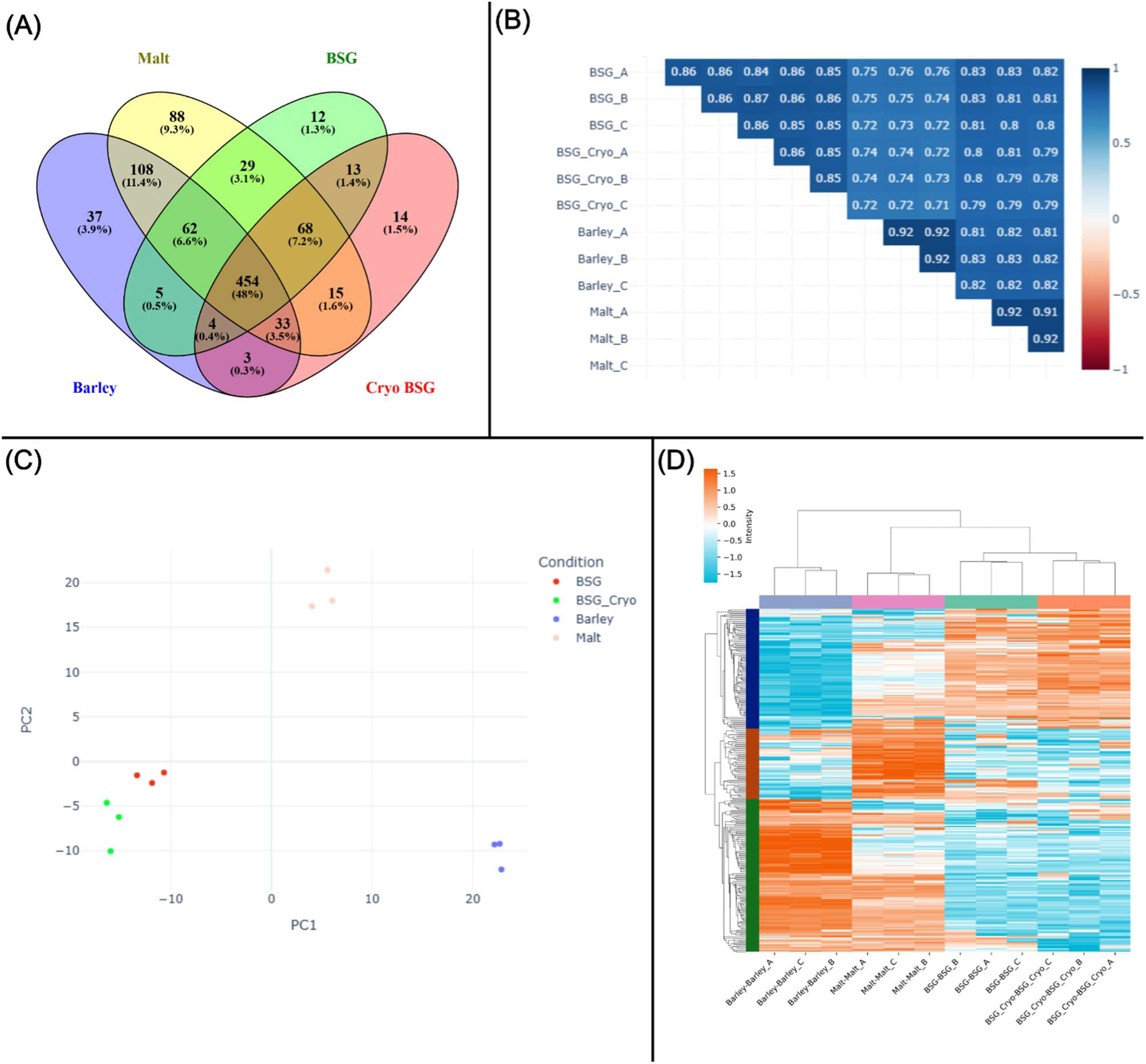
Overview of LC-MS/MS proteomics analysis. (A) Venn diagram showing overlap of reproducibly identified proteins (i.e. identified in at least two of three replicates) across the four samples showing both the numerical and relative (in %) intersect. (B) Pearson correlation matrix showing the overall qualitative and quantitative reproducibility between individual replicates and samples from anti-correlation (red, -1) over no correlation (white, 0) to perfect correlation (blue, 1).

Pearson correlation coefficients were based on MaxLFQ intensity data after imputation of missing values in MassDynamics. (C) Principal component analysis to show replicate-wise sample similarity and clustering according to overall qualitative and quantitative variability. Inidividual sample replicates are depicted based on the two first principal components describing together explaining 38% of the total variability (see Scree plot in Fig. S1). (D) Heatmap of differentially abundant proteins by ANOVA analysis in MassDynamics. Data is depicted as z-score standardized MaxLFQ intensities by row (protein group) and clustered using a Euclidian distance of 4.

The qualitative differences were also reflected when analyzing the overall replicate correlation (Fig. 3B) and experimental variability (Fig. 3C). Replicates between each sample displayed good overall correlation above 0.85 with grain samples generally showing higher replicate correlation (∼0.92). This difference may be ascribed to the higher number of protein identifications in these samples and a potentially more homogenous nature compared to BSG. Comparing BSG samples with and without cryo-milling showed comparable correlations as replicates within each group. This indicates minimal effect of the milling. As expected, BSG replicates had a substantially higher correlation with malt replicates than barley replicates, as the BSG was produced from the malt and because malting represent the biological germination process of the barley grains. These observations were also reflected when investigating the overall variability in the dataset using principal component analysis (PCA) (Fig. 3B). Here, the two BSG samples clustered nicely and have a longer distance (but in the same direction) from barley than does malt along the first principal component. In contrast, barley and BSG were closely spaced along PC2, while malt differs substantially. This indicates that differences between samples are governed by several underlying distributions, which was also corroborated by Scree analysis (Fig. S1), where PC1 and PC2 were found to explain only 24% and 14% of the total variability, respectively. Overall, this analysis nicely illustrates the dynamic process of moving from barley to BSG through malting and mashing. These dynamics are also evident when considering the 272 differentially expressed/abundant proteins by ANOVA across the different samples (28.8 % of reproducibly identified proteins), showing distinct clustering on the protein level (Fig. 3C).

### 3.4 Malting reduces abundance of stress-related proteins and induces enzymatic storage depletion

Malting of barley represents the biological process of germination. As such, this process is also expected to be reflected on the protein-level through significant changes in expression patterns. Through a differential expression analysis, we indeed found 27 proteins to be significantly (p < 0.05) downregulated (FC > 2) in malt while 69 proteins were found to be significantly upregulated based on normalized LFQ intensities (Fig. 4A). As the proteome is sub optimally annotated, we performed BLAST analysis against the Uniprot reference proteome. Overall, we found strong BLAST hits with identities above 90% (the majority at 100%) and e-values below 10^-50^ (Table S2). Among the 27 downregulated proteins in malt, we predominantly found proteins previously associated with germination. For instance, we found multiple late embryogenesis abundant (LEA) proteins, which have been shown to be involved with seed adaptation to particularly cold stress [65]. That these proteins are found to be depleted in the malted barley is not surprising as they have previously been shown to accumulate under certain stages of embryo development [66] and disappear after germination [67]. Similarly, other stress-related proteins such as the universal stress protein USP7364 were also found depleted in malt. Reduction in stress-related proteins may be considered a direct consequence of seeds transitioning from prolonged cold storage under dehydration to the germination phase, thereby entering a more normalized plant life cycle [68]. Several seed maturation proteins and fumarase, which are linked with dormancy seed maintenance and metabolism [69,70], were also found to be depleted. Furthermore, a large variety of depleted proteins are known to be involved in seed energy and nutrient storage such as subtilisin-chymotrypsin inhibitors, alpha-amylase inhibitors, hordeins, oleosin, and cupin-type embryo globulins [71–75]. Together, these findings clearly indicate how the seeds deactivate the dormant and inhibitory machinery and start consuming stored nutrients to enter the germination process.

**Figure 4:**
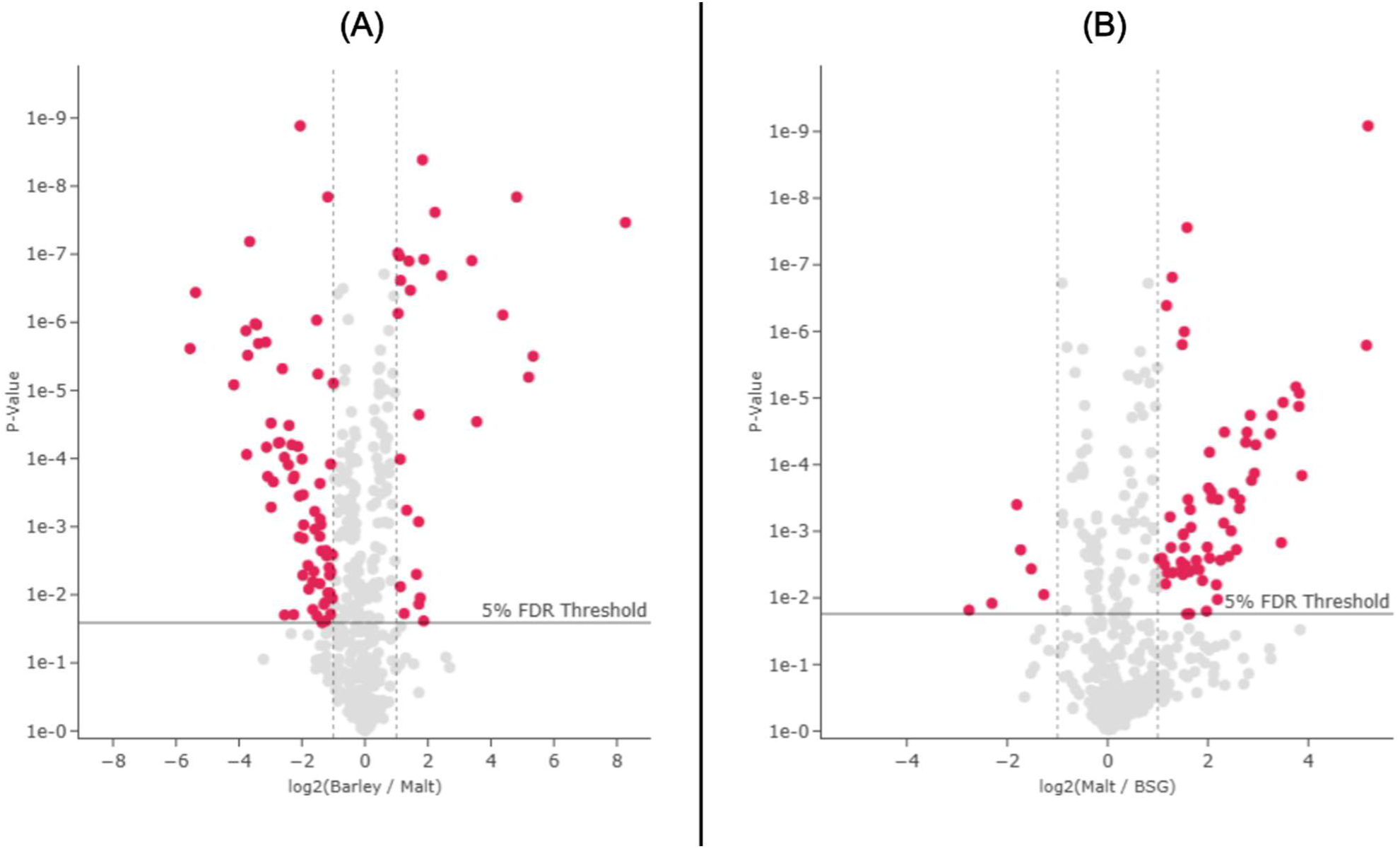
Volcano-plots from pairwise differential analysis of protein LFQ intensity for barley vs. malt (A) and malt vs. BSG (B). Highlighted proteins (in red) were found to be significantly (p < 0.05) and substantially (FC > 2) differentially abundant in the pairwise analysis in MassDynamics. An overview of differentially abundant proteins for barley vs. malt can be found in Table S3 while differentially abundant proteins from malt vs. BSG can be found in Table S4.

The upregulated subproteome in malt (69 proteins) is dominated by enzymes involved in various cellular pathways (Table S2). Not surprisingly, several isoforms of α-amylase were found to be upregulated. Additional carbohydrate degrading enzymes like β-mannosidase, α-glucosidase, β- glucanase, and chitinase, as well as other central enzymes in the carbohydrate metabolism such as malate synthase and Aldose 1-epimerase, were also found significantly (p < 0.05) and substantially (FC > 2) upregulated. Upregulation of carbohydrate degrading enzymes during malting has previously been demonstrated on both protein-[35] and RNA-level [76], as the seeds start to germinate and require liberation of stored nutrient. In addition to carbohydrate-related enzymes, a wide range of proteases were also found to be upregulated. This is also a direct consequence of the germination, where the seed storage proteins are degraded to facilitate initial plant growth [77]. This biological process is reflected in several other upregulated proteins such as lipoxygenases [78] and dehydrin, [79] as well as many other proteins involved in the central cellular machinery. In fact, the most upregulated protein is a histone, which further reflects the development of the seed embryo and initiation of growth.

The dramatic increase in enzymatic activity and initial plant development also induces increased oxidative stress on the seed [80]. This calls for protective measures to be activated against e.g. the increased formation of reactive oxygen species (ROS). Consequently, enzymes protecting against ROS, such as superoxide dismutase and peroxidase were also found to be upregulated during malting. Overall, these findings are in agreement with earlier studies on protein-level changes during barley seed germination and the malting process in brewing [41,81,82]. As such, this analysis functions as a built-in quality control of the analytical pipeline.

### 3.5 Selective enrichment and depletion during mashing and BSG isolation

Similarly to the pairwise analysis of barley and malt (Fig. 4A), significant protein-level differences were also found when comparing malt and BSG by LFQ intensities (Fig. 4B). Here, 66 proteins were found to be significantly (p < 0.05) depleted (FC > 2) in the BSG while only six were found to be significantly enriched. When comparing malt and BSG, it is important to refrain from the up-/downregulation terminology and rather describe enrichment and depletion, as mashing represents a physical processing of the malt, whereas malting represent a biological process within the barley seeds. As described in relation to investigating the malting process, we performed BLAST analysis for differential proteins found after mashing. Overall, we found strong BLAST hits with identities above 90% (the majority at 100%) and e-values below 10^-50^ (Table S3). Among the most substantially depleted proteins were infection defense-related such as hordoindoline-B1, α-hordothionin, and thaumatin as well as stress-related proteins such as Dehydrin, a LEA protein [83], and heat-shock proteins. The depletion of defense-related proteins may be considered a massive advantage for the use of BSG in foods, as e.g. thionin are linked with ubiquitous toxicity [84], while thaumatin is linked with increased allergenic risk [85]. Similarly, expansin [86] and non-specific lipid transfer proteins [87] are known allergens and were also depleted in BSG. Moreover, a range of different proteins identified as protease and amylase inhibitors, such as cystatin Hv-CPI5 and several bifunctional inhibitor/plant lipid transfer proteins, were depleted in BSG. This can be considered a highly beneficial change in the protein composition, as these inhibitors constitute anti-nutritional factors [88] and potential allergens [87], thereby increasing the safe applicability of BSG in foods. Of the six enriched proteins in BSG, only Glutathione S-transferase has previously been associated to allergy and has been classified as a minor allergen in birch pollen [89]. However, the abundance is below 0.05% (Table S1) and therefore not considered problematic as such being a minor allergen.

The selective depletion of proteins during mashing and subsequent removal of the wort is not only relevant on the protein-level but also on the bulk level. This is also illustrated from significant (p < 0.001) differences in the EAA/NEAA content between BSG and both barley and malt, while no significant difference was observed between the latter two (Fig. 1C). Moreover, this is also reflected when considering the cellular origin of the identified proteins considered as a whole (Fig. 5A). In barley, around 60% (mol/mol) of the total protein (by riBAQ) is predicted to be extracellular. This is significantly (p < 0.0001) reduced in both malt to 55 % but also significantly reduced (p < 0.0001) when comparing BSG (51%) to malt (Fig. S2). This indicates a continued depletion of the relative proportion of extracellular protein, while the relative share of cytoplasmic, nuclear, and plastid proteins significantly increase. Depletion of extracellular protein in general correlates well with the majority of depleted proteins identified from pairwise analysis (Fig. 4B) being of extracellular origin (Table SX). This indicates that extracellular proteins, to some extent, are either degraded during mashing or end up in the wort due to their highly accessible and soluble nature. This finding corroborates earlier studies showing the dynamic process of endogenous proteolysis during mashing [90] to which extracellular proteins would be more readily available.

**Figure 5:**
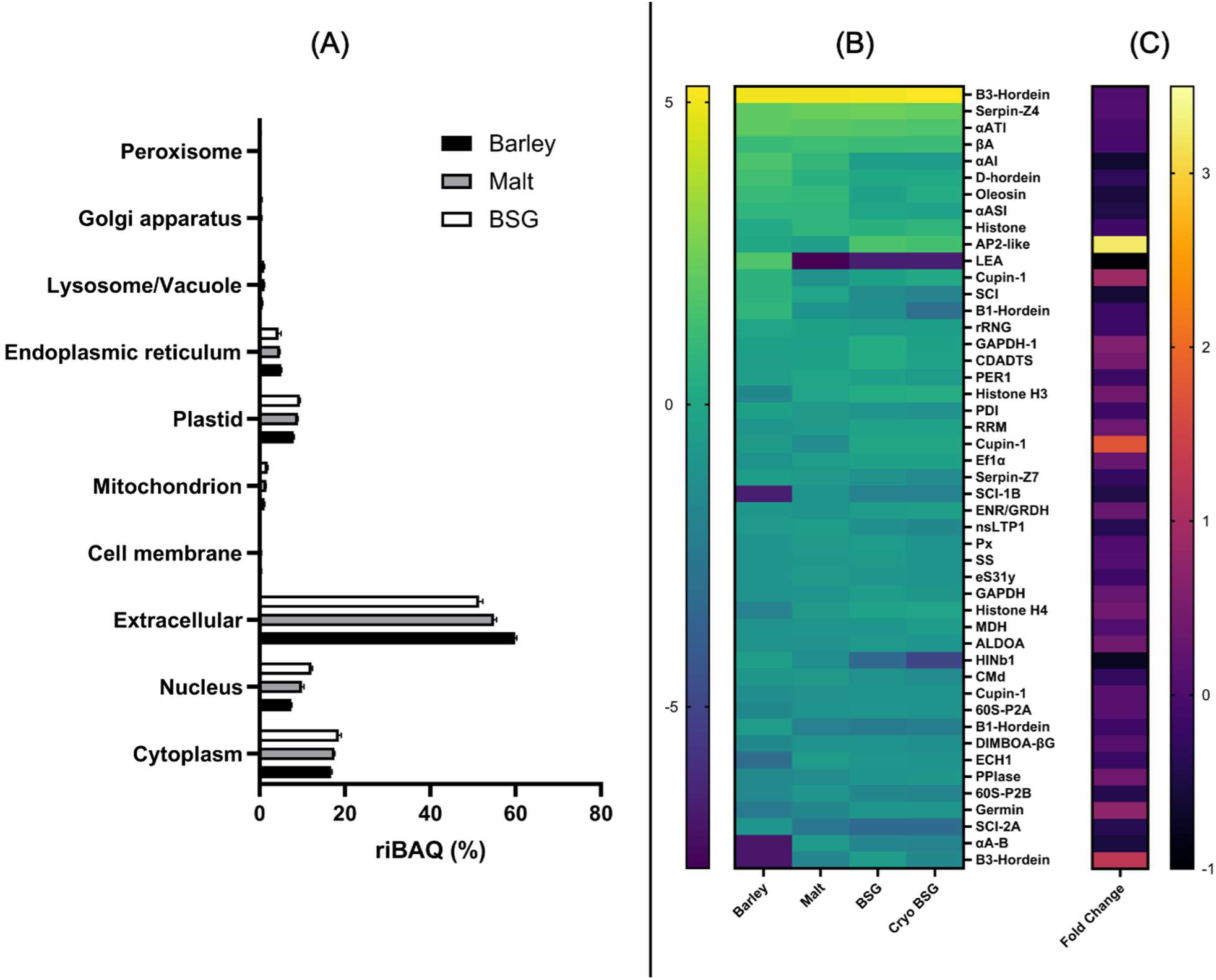
Relative molar abundance on subcellular and single protein level. (A) Distribution of subcellular localization as predicted by DeepLoc 2.0. Each compartment represents the sum riBAQ- based molar abundance for all proteins which has bas been ascribed hereto based on maximum probability. (B) Abundance distribution of the most abundant proteins across all samples (mean riBAQ > 0.5% in any sample). Protein abundance is depicted as the log2 transform of the relative molar abundance (by riBAQ [%]) and color coded from low (blue) to high (yellow) abundance. Proteins are given using short names as defined in Table S6. (C) Protein-level fold change (by riBAQ) of BSG vs. malt to visualize enrichment/depletion after mashing. Proteins are show from highly depleted (black) to highly enriched (yellow) in BSG and aligned with the protein names in panel B.

To obtain further insights and understanding hereof, it is relevant to consider absolute abundance changes on the single protein level over the full process. For this purpose, we determined the molar abundance at the single-protein level using riBAQ for proteins constituting a substantial amount of the total protein in any of the analyzed samples (Fig. 5B). The most abundant proteins (riBAQ > 0.5%) across all samples, B3-Hordein and Serpin-Z4, together constituting 36-43% of the total protein, show quite comparable levels of abundance in all samples. However, the LEA protein found abundant in barley (3.3%) was absent in downstream samples. This correlates with findings from the pairwise analysis of barley and malt (Fig. 4A). Several other proteins such as α-amylase inhibitor (αAI), D-hordein, oleosin, and subtilisin-chymotrypsin inhibitor (SCI) show a gradual decrease in abundance along the process and not only between individual process points, as previously described.

Interestingly, two proteins showed an opposite trend and became enriched and highly abundant in BSG. AP2-like factor was enriched threefold from barley (1.0% riBAQ) to BSG (3.0% riBAQ). This despite showing a minor depletion in malt (0.72% riBAQ). However, this protein was only identified from one peptide across all samples and received a low Andromeda score (6.1). Therefore, this protein identification is associated with uncertainty (Table S1). Cupin-type 1 embryo globulin had an abundance of 0.61% in barley, which decreased to 0.35% in malt but became enriched in BSG at 0.96%. These changes are also reflected by a fold change (FC) analysis between BSG and malt, where the two proteins obtained an FC of 3.2 and 1.8, respectively, based on molar abundance (Fig. 5C). Histone H3 showed a gradual increase from 0.31% in barley, over 0.86% in malt, to 1.2% in BSG. As such the BSG-to-malt FC is more subtle (0.39) but it may still be regarded as a highly abundant and enriched protein in BSG. The enrichment of a cupin-like protein may be disadvantageous as these are generally considered major allergens [91], and require additional processing to reduce allergenic potential of BSG as food ingredient.

When evaluating the effect of BSG cryo-milling, we found that this did not entail an overall positive effect as anticipated. Firstly, there was no real difference in the protein profile by SDS-PAGE when compared to the crude BSG (Fig. 1B, 2B). Moreover, the number of reproducibly identified proteins actually dropped by 7% from 647 in the crude BSG to 604 in the cryo-milled BSG. While the overall differences from differential analysis by ANOVA (Fig. 3D) and pairwise analysis (Fig. S3) were minimal, the overall loss of protein identifications and introduction of variance between BSG and cryo-milled BSG (Fig. 3C, 5B) indicates that the cryo-milling did not improve the analytical output.

In fact, the additional processing steps may have introduced selective losses, which is why the analytical outcome may not fully reflect the BSG and could be considered somewhat detrimental instead. The fact that no additional improvement was obtained from BSG cryo-milling may be a result of the BSG already being quite heavily processed and can be considered equivalent to a residual grain flour. This finding facilitates easier and faster sample preparation for BSG proteomics analysis.

3.6 Highly abundant proteins in BSG: Risk factors or potential targets for downstream processing?

With the aim of identifying highly abundant proteins that could be potential targets for downstream processing and BSG valorization, we further investigated the most abundant proteins (> 1% riBAQ) in BSG (Table 1). While BSG was found to be depleted in many proteins with antinutritional and allergenic potential, it is also clear that several of the most abundant proteins in BSG are still associated with risk factors for safe ingestion in foods. In a previous proteomic study of BSG [92], many of the same proteins were identified as abundant. However, this study quantified protein abundance by relative LFQ intensity, thereby normalizing data between samples and not within the individual samples as done with riBAQ. This also entails that the relative protein distribution in that study did not reflect the molar distribution but rather the relative signal distribution, where the abundance of larger proteins was artificially inflated. Nevertheless, the MW of the most abundant proteins, in particular B3-Hordein, Serpin-Z4, and alpha-amylase/trypsin inhibitor (αATI), corresponded well with bands observed by SDS-PAGE analysis of BSG (Fig. 1B, 2B). This indicates that these proteins remained intact after mashing, while the substantial increase in a diffuse smear in the low MW region of BSG compared to both barely and malt, could indicate that some proteins may be partially digested in the process.

**Table 1:**
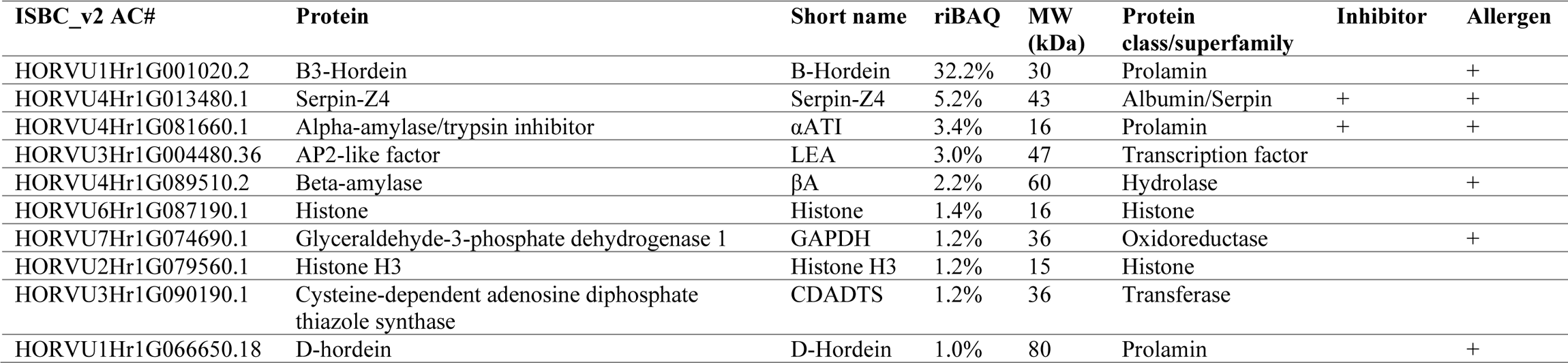
Overview of abundant (> 1%) BSG proteins by relative molar abundance (riBAQ). Table shows the ISBC_v2 AC#, the protein name (as inferred by BLAST analysis in Uniprot), short name, mean riBAQ (%) in BSG, molecular weight (MW) in kDa, the class and/or protein superfamily (if relevant), and know association with inhibitory potential against protein and/or carbohydrate digestive enzymes as well as allergenic potential based on protein superfamily or homology as indicated in literature (indicated by “+”).

Hordeins, including the abundant B-Hordein and D-Hordein, are prolamins and thereby insoluble [93]. In addition to their high natural abundance in barley, their insolubility makes it logical that residual hordeins end up in the solid BSG fraction after mashing. Hordeins are homologues to wheat gluten proteins (gliadins/glutelins), and hence also considered major allergens [91]. Being rich in glutamine and proline [94], the high hordein content in not only BSG but all analyzed samples are a major driver of the high content of these AAs. Serpin-Z4 is considered a part of the albumin superfamily and is known to elicit serine protease inhibition. As such it is classified as an antinutritional factor [95]. Earlier studies found that serpin-Z4 survives the entire brewing process and is in fact positively correlated with beer foaming and considered a marker of foam stability [96]. Moreover, Serpin-Z4 was also found to survive gastrointestinal digestion in ruminants [95], which further highlights its limited direct nutritional value. Similar results were found for αATI [95]. Both Serpins and αATIs are known grain allergens [91,97]. In addition, Glyceraldehyde-3-phosphate dehydrogenase 1 (GADPH) is a known wheat allergen affiliated with Baker’s Asthma [98]. For the remaining highly abundant proteins, we were only able to find literature on their specific cellular function but no reports of cytotoxicity, antinutritional effect, or allergenic potential.

Ultimately, and based on only the most abundant proteins, more than 45% of the BSG proteins are either associated with allergenicity or considered antinutritional. This further substantiates the direct use of BSG may be complicated in terms of food safety and nutritional value. As such, further processing is required to increase BSG utilization in a food context. While the major proteins of BSG may be considered potential risk factors for its application in foods, targeting these proteins for downstream processing by e.g. hydrolysis should be further explored.

## 4. Conclusions

From the enormous global beer production, BSG constitutes the largest side-stream with an annual production of around 40 MT. While substantial effort is put toward the efficient use of this by-product, the current developments still make the side-stream a largely underutilized waste-stream. By focusing on the molecular makeup of the BSG, it may be possible to develop processes for valorization of BSG facilitating better resource utilization. In this study, we performed a protein-centric comparison of barley, malt, and BSG to not only provide a detailed characterization of the BSG but also how this protein composition originates. BSG was found to have a significantly higher protein content (17.4%) than barley and malt (9.0% and 10.5%, respectively), albeit still modest in proportion for use as a protein-based food ingredient. Through bottom-up proteomics, we identified very distinct and significant protein-level changes between the three stages of the brewing process. Malting was found to reduce levels of stress related proteins while facilitating increased nutrient use by reducing endogenous enzyme inhibitors and enriching active enzymes. As a result of the mashing process and subsequent wort removal, the BSG was found to be depleted in a wide range of proteins also covering additional stress-related proteins and amylase/protease inhibitors. This finding corelated with an overall reduction of proteins of extracellular origin in BSG. While this reduction is beneficial in terms of food safety, a large proportion of BSG (> 45%) is still constituted by proteins with known antinutritional and allergenic properties, whereof hordeins represent the majority. Nevertheless, BSG was shown to have a good amino acid profile, with only Lys falling slightly below the recommended proportion of essential amino acids. As such, BSG can be considered a high-potential source of functional protein-based ingredients, but it requires development and adaptation of adequate technology to ensure the production of a protein-rich and safe food ingredient.

## Data availability

The mass spectrometry proteomics data have been deposited into the ProteomeXchange Consortium via the PRIDE [49] partner repository with the dataset identifier PXD062029 and 10.6019/PXD062029. Numerical data for crude characterization as well as MaxQuant output data and downstream analysis thereof is available in the supplementary material.

## Author contributions

SGE: Conceptualization, Methodology, Validation, Formal analysis, Investigation, Writing – original draft preparation, Writing – review and editing, Visualization, Funding acquisition, Project administration, Supervision. RKM: Methodology, Validation, Formal analysis, Investigation, Writing – original draft preparation, Writing – review and editing. NAK: Methodology, Validation, Formal analysis, Investigation, Writing – review and editing. LSQ: Methodology, Validation, Writing – review and editing. TJH: Conceptualization, Methodology, Writing – review and editing, Funding acquisition, Supervision. BS: Conceptualization, Writing – review and editing. MTO: Conceptualization, Writing – review and editing, Funding acquisition. CJ: Conceptualization, Writing – review and editing, Funding acquisition, Project administration, Supervision. BY: Conceptualization, Writing – review and editing, Funding acquisition, Project administration.

All authors have read and agreed to the published version of the manuscript.

## Acknowledgements

The authors would like to acknowledge Viking Malt (Vordingborg, Denmark) for supply of barley and malt used in this work. The authors would also like to thank Inge Houlberg Holmberg for her assistance with amino acid analysis.

## References

1 Devnani, B., Moran, G.C. and Grossmann, L. (2023) Extraction, Composition, Functionality, and Utilization of Brewer’s Spent Grain Protein in Food Formulations. Foods 2023, Vol. 12, Page 1543, Multidisciplinary Digital Publishing Institute, 12, 1543. 10.3390/FOODS12071543.

2 Hejna, A. (2021) More than Just a Beer—the Potential Applications of by-Products from Beer Manufacturing in Polymer Technology. Emergent Materials 2021 5:3, Springer, 5, 765–783. 10.1007/S42247-021-00304-4.

3 Mussatto, S.I. (2014) Brewer’s Spent Grain: A Valuable Feedstock for Industrial Applications. Journal of the Science of Food and Agriculture, John Wiley & Sons, Ltd, 94, 1264–1275. 10.1002/JSFA.6486.

4 Bjerregaard, M.F., Charalampidis, A., Frøding, R., Shetty, R., Pastell, H., Jacobsen, C., Zhuang, S., Pinelo, M., Hansen, P.B. and Hobley, T.J. (2019) Processing of Brewing By- Products to Give Food Ingredient Streams. European Food Research and Technology, Springer Verlag, 245, 545–558. 10.1007/S00217-018-03224-6/FIGURES/5.

5 Wen, C., Zhang, J., Duan, Y., Zhang, H. and Ma, H. (2019) A Mini-Review on Brewer’s Spent Grain Protein: Isolation, Physicochemical Properties, Application of Protein, and Functional Properties of Hydrolysates. Journal of Food Science, John Wiley & Sons, Ltd, 84, 3330–3340. 10.1111/1750-3841.14906.

6 Nyhan, L., Sahin, A.W., Schmitz, H.H., Siegel, J.B. and Arendt, E.K. (2023) Brewers’ Spent Grain: An Unprecedented Opportunity to Develop Sustainable Plant-Based Nutrition Ingredients Addressing Global Malnutrition Challenges. Journal of Agricultural and Food Chemistry, American Chemical Society, 71, 10543–10564. 10.1021/ACS.JAFC.3C02489/ASSET/IMAGES/LARGE/JF3C02489_0005.JPEG.

7 Lynch, K.M., Steffen, E.J. and Arendt, E.K. (2016) Brewers’ Spent Grain: A Review with an Emphasis on Food and Health. Journal of the Institute of Brewing, John Wiley & Sons, Ltd, 122, 553–568. 10.1002/JIB.363.

8 Ikram, S., Huang, L.Y., Zhang, H., Wang, J. and Yin, M. (2017) Composition and Nutrient Value Proposition of Brewers Spent Grain. Journal of Food Science, John Wiley & Sons, Ltd, 82, 2232–2242. 10.1111/1750-3841.13794.

9 Steiner, J., Procopio, S. and Becker, T. (2015) Brewer’s Spent Grain: Source of Value-Added Polysaccharides for the Food Industry in Reference to the Health Claims. European Food Research and Technology, Springer Verlag, 241, 303–315. 10.1007/S00217-015-2461-7/TABLES/3.

10 Vieira, E., Rocha, M.A.M., Coelho, E., Pinho, O., Saraiva, J.A., Ferreira, I.M.P.L.V.O. and Coimbra, M.A. (2014) Valuation of Brewer’s Spent Grain Using a Fully Recyclable Integrated Process for Extraction of Proteins and Arabinoxylans. Industrial Crops and Products, Elsevier, 52, 136–143. 10.1016/J.INDCROP.2013.10.012.

11 Connolly, A., Piggott, C.O. and Fitzgerald, R.J. (2013) Characterisation of Protein-Rich Isolates and Antioxidative Phenolic Extracts from Pale and Black Brewers’ Spent Grain. International Journal of Food Science and Technology, Oxford Academic, 48, 1670–1681. 10.1111/IJFS.12137.

12 Qin, F., Johansen, A.Z. and Mussatto, S.I. (2018) Evaluation of Different Pretreatment Strategies for Protein Extraction from Brewer’s Spent Grains. Industrial Crops and Products, Elsevier, 125, 443–453. 10.1016/J.INDCROP.2018.09.017.

13 Rommi, K., Niemi, P., Kemppainen, K. and Kruus, K. (2018) Impact of Thermochemical Pre-Treatment and Carbohydrate and Protein Hydrolyzing Enzyme Treatment on Fractionation of Protein and Lignin from Brewer’s Spent Grain. Journal of Cereal Science, Academic Press, 79, 168–173. 10.1016/J.JCS.2017.10.005.

14 Niemi, P., Martins, D., Buchert, J. and Faulds, C.B. (2013) Pre-Hydrolysis with Carbohydrases Facilitates the Release of Protein from Brewer’s Spent Grain. Bioresource Technology, Elsevier, 136, 529–534. 10.1016/J.BIORTECH.2013.03.076.

15. Abeynayake, R., Zhang, S., Yang, W. and Chen, L. (2022) Development of Antioxidant Peptides from Brewers’ Spent Grain Proteins. LWT, Academic Press, 158, 113162. 10.1016/J.LWT.2022.113162.

16 Yalçın, E., Çelik, S. and İbanoğlu, E. (2008) Foaming Properties of Barley Protein Isolates and Hydrolysates. European Food Research and Technology, Springer Verlag, 226, 967– 974. 10.1007/S00217-007-0618-8/FIGURES/10.

17 Yalçın, E., Çelik, S. and İbanoğlu, E. (2008) Foaming Properties of Barley Protein Isolates and Hydrolysates. European Food Research and Technology, Springer Verlag, 226, 967– 974. 10.1007/S00217-007-0618-8/FIGURES/10.

18 Xia, Y., Bamdad, F., Gänzle, M. and Chen, L. (2012) Fractionation and Characterization of Antioxidant Peptides Derived from Barley Glutelin by Enzymatic Hydrolysis. Food Chemistry, Elsevier, 134, 1509–1518. 10.1016/J.FOODCHEM.2012.03.063.

19 Ikram, S., Zhang, H., Ahmed, M.S. and Wang, J. (2020) Ultrasonic Pretreatment Improved the Antioxidant Potential of Enzymatic Protein Hydrolysates from Highland Barley Brewer’s Spent Grain (BSG). Journal of Food Science, John Wiley & Sons, Ltd, 85, 1045–1059. 10.1111/1750-3841.15063.

20 Olsen, T.H., Yesiltas, B., Marin, F.I., Pertseva, M., García-Moreno, P.J., Gregersen, S., Overgaard, M.T., Jacobsen, C., Lund, O., Hansen, E.B. and Marcatili, P. (2020) AnOxPePred: Using Deep Learning for the Prediction of Antioxidative Properties of Peptides. Scientific Reports, Nature Research, 10, 21471. 10.1038/s41598-020-78319-w.

21 García-Moreno, P.J., Jacobsen, C., Marcatili, P., Gregersen, S., Overgaard, M.T., Andersen, M.L., Sørensen, A.D.M. and Hansen, E.B. (2020) Emulsifying Peptides from Potato Protein Predicted by Bioinformatics: Stabilization of Fish Oil-in-Water Emulsions. Food Hydrocolloids, Elsevier B.V., 101, 105529. 10.1016/j.foodhyd.2019.105529.

22 García-Moreno, P.J., Gregersen, S., Nedamani, E.R., Olsen, T.H., Marcatili, P., Overgaard, M.T., Andersen, M.L., Hansen, E.B. and Jacobsen, C. (2020) Identification of Emulsifier Potato Peptides by Bioinformatics: Application to Omega-3 Delivery Emulsions and Release from Potato Industry Side Streams. Scientific Reports, Nature Research, 10, 690. 10.1038/s41598-019-57229-6.

23 García-Moreno, P.J., Yesiltas, B., Gregersen Echers, S., Marcatili, P., Overgaard, M.T., Hansen, E.B. and Jacobsen, C. (2023) Recent Advances in the Production of Emulsifying Peptides with the Aid of Proteomics and Bioinformatics. Current Opinion in Food Science, Elsevier, 51, 101039. 10.1016/J.COFS.2023.101039.

24 Varona, E., García-Moreno, P.J., Gregersen Echers, S., Olsen, T.H., Marcatili, P., Guardiola, F., Overgaard, M.T., Hansen, E.B., Jacobsen, C. and Yesiltas, B. (2023) Antioxidant Peptides from Alternative Sources Reduce Lipid Oxidation in 5% Fish Oil-in-Water Emulsions (PH 4) and Fish Oil-Enriched Mayonnaise. Food Chemistry, Elsevier, 426, 136498. 10.1016/J.FOODCHEM.2023.136498.

25 Yesiltas, B., Soria Caindec, A.M., García-Moreno, P.J., Echers, S.G., Olsen, T.H., Jones, N.C., Hoffmann, S. V., Marcatili, P., Overgaard, M.T., Hansen, E.B. and Jacobsen, C. (2023) Physical and Oxidative Stability of Fish Oil-in-Water Emulsions Stabilized with Emulsifier Peptides Derived from Seaweed, Methanotrophic Bacteria and Potato Proteins. Colloids and Surfaces A: Physicochemical and Engineering Aspects, Elsevier, 663, 131069. 10.1016/J.COLSURFA.2023.131069.

26 Yesiltas, B., García-Moreno, P.J., Gregersen, S., Olsen, T.H., Jones, N.C., Hoffmann, S. V., Marcatili, P., Overgaard, M.T., Hansen, E.B. and Jacobsen, C. (2022) Antioxidant Peptides Derived from Potato, Seaweed, Microbial and Spinach Proteins: Oxidative Stability of 5% Fish Oil-in-Water Emulsions. Food Chemistry, Elsevier, 385, 132699. 10.1016/J.FOODCHEM.2022.132699.

27 García-Moreno, P.J., Yang, J., Gregersen, S., Jones, N.C., Berton-Carabin, C.C., Sagis, L.M.C., Hoffmann, S. V., Marcatili, P., Overgaard, M.T., Hansen, E.B. and Jacobsen, C. (2021) The Structure, Viscoelasticity and Charge of Potato Peptides Adsorbed at the Oil- Water Interface Determine the Physicochemical Stability of Fish Oil-in-Water Emulsions. Food Hydrocolloids, Elsevier, 115, 106605. https://linkinghub.elsevier.com/retrieve/pii/S0268005X21000217.

28 Gregersen Echers, S., Abdul-Khalek, N., Mikkelsen, R.K., Holdt, S.L., Jacobsen, C., Hansen, E.B., Olsen, T.H., Sejberg, J.J.P. and Overgaard, M.T. (2022) Is Gigartina a Potential Source of Food Protein and Functional Peptide-Based Ingredients? Evaluating an Industrial, Pilot- Scale Extract by Proteomics and Bioinformatics. Future Foods, Elsevier, 6, 100189. 10.1016/J.FUFO.2022.100189.

29 Yesiltas, B., Gregersen, S., Lægsgaard, L., Brinch, M.L., Olsen, T.H., Marcatili, P., Overgaard, M.T., Hansen, E.B., Jacobsen, C. and García-Moreno, P.J. (2021) Emulsifier Peptides Derived from Seaweed, Methanotrophic Bacteria, and Potato Proteins Identified by Quantitative Proteomics and Bioinformatics. Food Chemistry, Elsevier, 362, 130217. 10.1016/J.FOODCHEM.2021.130217.

30 Deka, A. and Saikia, V. (2023) Prediction of ACE-Inhibitory and Antioxidant Peptides from Storage Proteins of Edible Seeds and Nuts Using in Silico Approaches. Sustainable Food Proteins, John Wiley & Sons, Ltd, 1, 84–98. 10.1002/SFP2.1013.

31 Gregersen Echers, S., Jafarpour, A., Yesiltas, B., García-Moreno, P.J., Greve-Poulsen, M., Hansen, D.K., Jacobsen, C., Overgaard, M.T. and Hansen, E.B. (2023) Targeted Hydrolysis of Native Potato Protein: A Novel Workflow for Obtaining Hydrolysates with Improved Interfacial Properties. Food Hydrocolloids, Elsevier, 137, 108299. 10.1016/J.FOODHYD.2022.108299.

32 Bjørlie, M., Yesiltas, B., García-Moreno, P.J., Espejo-Carpio, F.J., Rahmani-Manglano, N.E., Guadix, E.M., Jafarpour, A., Hansen, E.B., Marcatili, P., Overgaard, M.T., Gregersen Echers, S. and Jacobsen, C. (2023) Bioinformatically Predicted Emulsifying Peptides and Potato Protein Hydrolysate Improves the Oxidative Stability of Microencapsulated Fish Oil. Food Chemistry Advances, Elsevier, 3, 100441. 10.1016/J.FOCHA.2023.100441.

33 Schulz, B.L., Phung, T.K., Bruschi, M., Janusz, A., Stewart, J., Meehan, J., Healy, P., Nouwens, A.S., Fox, G.P. and Vickers, C.E. (2018) Process Proteomics of Beer Reveals a Dynamic Proteome with Extensive Modifications. Journal of Proteome Research, American Chemical Society, 17, 1647–1653. 10.1021/ACS.JPROTEOME.7B00907/ASSET/IMAGES/LARGE/PR-2017-009078_0006.JPEG.

34 Blanco, C.A., Caballero, I., Barrios, R. and Rojas, A. (2014) Innovations in the Brewing Industry: Light Beer. International Journal of Food Sciences and Nutrition, Informa UK Ltd, 65, 655–660. 10.3109/09637486.2014.893285.

35 Bahmani, M., O’Lone, C.E., Juhász, A., Nye-Wood, M., Dunn, H., Edwards, I.B. and Colgrave, M.L. (2021) Application of Mass Spectrometry-Based Proteomics to Barley Research. Journal of Agricultural and Food Chemistry, American Chemical Society, 69, 8591–8609. 10.1021/ACS.JAFC.1C01871/ASSET/IMAGES/LARGE/JF1C01871_0002.J PEG.

36 Kerr, E.D., Fox, G.P. and Schulz, B.L. (2023) Proteomics and Metabolomics Reveal That an Abundant α-Glucosidase Drives Sorghum Fermentability for Beer Brewing. Journal of Proteome Research, American Chemical Society, 22, 3596–3606. 10.1021/ACS.JPROTEOME.3C00436/ASSET/IMAGES/LARGE/PR3C00436_0006.JPEG.

37 Mahalingam, R. (2018) Temporal Analyses of Barley Malting Stages Using Shotgun Proteomics. PROTEOMICS, John Wiley & Sons, Ltd, 18, 1800025. 10.1002/PMIC.201800025.

38 Liu, S., Kerr, E.D., Pegg, C.L. and Schulz, B.L. (2022) Proteomics and Glycoproteomics of Beer and Wine. PROTEOMICS, John Wiley & Sons, Ltd, 22, 2100329. 10.1002/PMIC.202100329.

39 Colgrave, M.L., Goswami, H., Howitt, C.A. and Tanner, G.J. (2013) Proteomics as a Tool to Understand the Complexity of Beer. Food Research International, Elsevier, 54, 1001–1012. 10.1016/J.FOODRES.2012.09.043.

40 Nájera-Torres, E., Bernal-Gracida, L.A., González-Solís, A., Schulte-Sasse, M., Castañón- Suárez, C., Juárez-Díaz, J.A., Cruz-Zamora, Y., Vázquez-Santana, S., Figueroa, M. and Cruz-García, F. (2022) Proteolytic Activities and Profiles as Useful Traits to Select Barley Cultivars for Beer Production. Journal of Food Biochemistry, John Wiley & Sons, Ltd, 46, e14094. 10.1111/JFBC.14094.

41 Qin, Q., Liu, J., Hu, S., Dong, J., Yu, J., Fang, L., Huang, S. and Wang, L. (2021) Comparative Proteomic Analysis of Different Barley Cultivars during Seed Germination. Journal of Cereal Science, Academic Press, 102, 103357. 10.1016/J.JCS.2021.103357.

42 Jin, Z., Cai, G.L., Li, X.M., Gao, F., Yang, J.J., Lu, J. and Dong, J.J. (2014) Comparative Proteomic Analysis of Green Malts between Barley (Hordeum Vulgare) Cultivars. Food Chemistry, Elsevier, 151, 266–270. 10.1016/J.FOODCHEM.2013.11.065.

43 Mikkelsen, R.K., Queiroz, L., Echers, S.G., Hobley, T., Overgaard, M., Jacobsen, C. and Svensson, B. (2024) Extracting Proteins from Brewers’ Spent Grain Using Emerging Technologies: Evaluating Efficiency and Use as Emulsifier. Authorea Preprints, Authorea. 10.22541/AU.173142929.94999881/V1.

44 Ghelichi, S., Sørensen, A.D.M., Hajfathalian, M. and Jacobsen, C. (2024) Effect of Post- Extraction Ultrasonication on Compositional Features and Antioxidant Activities of Enzymatic/Alkaline Extracts of Palmaria Palmata. Marine Drugs 2024, Vol. 22, Page 179, Multidisciplinary Digital Publishing Institute, 22, 179. 10.3390/MD22040179.

45 Gregersen, S., Kongsted, A.-S.H., Nielsen, R.B., Hansen, S.S., Lau, F.A., Rasmussen, J.B., Holdt, S.L. and Jacobsen, C. (2021) Enzymatic Extraction Improves Intracellular Protein Recovery from the Industrial Carrageenan Seaweed Eucheuma Denticulatum Revealed by Quantitative, Subcellular Protein Profiling: A High Potential Source of Functional Food Ingredients. Food Chemistry: X, 12, 100137. 10.1016/J.FOCHX.2021.100137.

46. 46 Tyanova, S., Temu, T. and Cox, J. (2016) The MaxQuant Computational Platform for Mass Spectrometry-Based Shotgun Proteomics. Nature Protocols, Nature Publishing Group, 11, 2301–2319. 10.1038/nprot.2016.136.

47 Kersey, P.J., Allen, J.E., Allot, A., Barba, M., Boddu, S., Bolt, B.J., Carvalho-Silva, D., Christensen, M., Davis, P., Grabmueller, C., Kumar, N., Liu, Z., Maurel, T., Moore, B., McDowall, M.D., Maheswari, U., Naamati, G., Newman, V., Ong, C.K., Paulini, M., Pedro, H., Perry, E., Russell, M., Sparrow, H., Tapanari, E., Taylor, K., Vullo, A., Williams, G., Zadissia, A., Olson, A., Stein, J., Wei, S., Tello-Ruiz, M., Ware, D., Luciani, A., Potter, S., Finn, R.D., Urban, M., Hammond-Kosack, K.E., Bolser, D.M., De Silva, N., Howe, K.L., Langridge, N., Maslen, G., Staines, D.M. and Yates, A. (2018) Ensembl Genomes 2018: An Integrated Omics Infrastructure for Non-Vertebrate Species. Nucleic Acids Research, Oxford Academic, 46, D802–D808. 10.1093/NAR/GKX1011.

48 Lee, S., Lee, T., Yang, S. and Lee, I. (2020) BarleyNet: A Network-Based Functional Omics Analysis Server for Cultivated Barley, Hordeum Vulgare L. Frontiers in Plant Science, Frontiers Media S.A., 11, 510185. 10.3389/FPLS.2020.00098/BIBTEX.

49 Perez-Riverol, Y., Bai, J., Bandla, C., García-Seisdedos, D., Hewapathirana, S., Kamatchinathan, S., Kundu, D.J., Prakash, A., Frericks-Zipper, A., Eisenacher, M., Walzer, M., Wang, S., Brazma, A. and Vizcaíno, J.A. (2022) The PRIDE Database Resources in 2022: A Hub for Mass Spectrometry-Based Proteomics Evidences. Nucleic Acids Research, Oxford Academic, 50, D543–D552. 10.1093/NAR/GKAB1038.

50 The UniProt Consortium. (2017) UniProt: The Universal Protein Knowledgebase. Nucleic Acids Research, 45, D158–D169. 10.1093/nar/gkw1099.

51 Cox, J., Hein, M.Y., Luber, C.A., Paron, I., Nagaraj, N. and Mann, M. (2014) Accurate Proteome-Wide Label-Free Quantification by Delayed Normalization and Maximal Peptide Ratio Extraction, Termed MaxLFQ. Molecular and Cellular Proteomics, American Society for Biochemistry and Molecular Biology Inc., 13, 2513–2526. 10.1074/MCP.M113.031591/ATTACHMENT/B414BAF4-20AE-46D2-BB85-A06FBF3A11C3/MMC1.ZIP.

52 Quaglieri, A., Bloom, J., Triantafyllidis, A., Green, B., Condina, M.R., Ngov, P.B., Infusini, G. and Webb, A.I. (2022) Mass Dynamics 2.0: An Improved Modular Web-Based Platform for Accelerated Proteomics Insight Generation and Decision Making. bioRxiv, Cold Spring Harbor Laboratory, 2022.12.12.517480. 10.1101/2022.12.12.517480.

53. 53 Schwanhüusser, B., Busse, D., Li, N., Dittmar, G., Schuchhardt, J., Wolf, J., Chen, W. and Selbach, M. (2011) Global Quantification of Mammalian Gene Expression Control. Nature, Nature Publishing Group, 473, 337–342. 10.1038/nature10098.

54 Danner Aakjaer Pedersen, K., Thopholm Andersen, L., Heiselberg, M., Agerskov Brigsted, C., Lyngs Støvring, F., Mailund Mikkelsen, L., Albrekt Hansen, S., Enrico Rusbjerg-Weberskov, C., Lübeck, M. and Gregersen Echers, S. (2025) Identifying Endogenous Proteins of Perennial Ryegrass (Lolium Perenne) with Ex Vivo Antioxidant Activity. Proteomes 2025, Vol. 13, Page 8, Multidisciplinary Digital Publishing Institute, **13**, 8. 10.3390/PROTEOMES13010008.

55 Thumuluri, V., Almagro Armenteros, J.J., Johansen, A.R., Nielsen, H. and Winther, O. (2022) DeepLoc 2.0: Multi-Label Subcellular Localization Prediction Using Protein Language Models. Nucleic Acids Research, Oxford Academic, 50, W228–W234. 10.1093/NAR/GKAC278.

56 Karlsen, F. and Skov, P. V. (2022) Review – Potentials and Limitations of Utilising Brewer’s Spent Grain as a Protein Source in Aquaculture Feeds. Journal of Cleaner Production, Elsevier, 357, 131986. 10.1016/J.JCLEPRO.2022.131986.

57 Jaeger, A., Zannini, E., Sahin, A.W. and Arendt, E.K. (2021) Barley Protein Properties, Extraction and Applications, with a Focus on Brewers’ Spent Grain Protein. Foods 2021, Vol. 10, Page 1389, Multidisciplinary Digital Publishing Institute, 10, 1389. 10.3390/FOODS10061389.

58 Bi, X., Ye, L., Lau, A., Kok, Y.J., Zheng, L., Ng, D., Tan, K., Ow, D., Ananta, E., Vafiadi, C. and Muller, J. (2018) Proteomic Profiling of Barley Spent Grains Guides Enzymatic Solubilization of the Remaining Proteins. Applied Microbiology and Biotechnology, Springer Verlag, 102, 4159–4170. 10.1007/S00253-018-8886-8/FIGURES/3.

59 Celus, I., Brijs, K. and Delcour, J.A. (2006) The Effects of Malting and Mashing on Barley Protein Extractability. Journal of Cereal Science, Academic Press, 44, 203–211. 10.1016/J.JCS.2006.06.003.

60. Gorissen, S.H.M., Crombag, J.J.R., Senden, J.M.G., Waterval, W.A.H., Bierau, J., Verdijk, L.B. and van Loon, L.J.C. (2018) Protein Content and Amino Acid Composition of Commercially Available Plant-Based Protein Isolates. Amino Acids, Springer-Verlag Wien, 50, 1685–1695. 10.1007/S00726-018-2640-5/TABLES/2.

61. WHO/FAO/UNU. (2007) PROTEIN AND AMINO ACID REQUIREMENTS IN HUMAN NUTRITION. World Healh Organization, Geneva. https://apps.who.int/iris/bitstream/handle/10665/43411/WHO_TRS_935_eng.pdf?sequence=1&isAllowed=y.

62 Cun, Z., Li, X., Zhang, J.Y., Hong, J., Gao, L.L., Yang, J., Ma, S.Y. and Chen, J.W. (2024) Identification of Candidate Genes and Residues for Improving Nitrogen Use Efficiency in the N-Sensitive Medicinal Plant Panax Notoginseng. BMC Plant Biology, BioMed Central Ltd, 24, 1–20. 10.1186/S12870-024-04768-4/FIGURES/12.

63 Berni, R., Leclercq, C.C., Roux, P., Hausman, J.F., Renaut, J. and Guerriero, G. (2023) A Molecular Study of Italian Ryegrass Grown on Martian Regolith Simulant. Science of The Total Environment, Elsevier, 854, 158774. 10.1016/J.SCITOTENV.2022.158774.

64 Lin, Z., Schaefer, K., Lui, I., Yao, Z., Fossati, A., Swaney, D.L., Palar, A., Sali, A. and Wells, J.A. (2024) Multiscale Photocatalytic Proximity Labeling Reveals Cell Surface Neighbors on and between Cells. Science (New York, N.Y.), American Association for the Advancement of Science, 385, eadl5763. 10.1126/SCIENCE.ADL5763/SUPPL_FILE/SCIENCE.ADL5763_MDAR_REPRODUCIBILITY_CHECKLIST.PDF.

65 Zan, T., Li, L., Li, J., Zhang, L. and Li, X. (2020) Genome-Wide Identification and Characterization of Late Embryogenesis Abundant Protein-Encoding Gene Family in Wheat: Evolution and Expression Profiles during Development and Stress. Gene, Elsevier, 736, 144422. 10.1016/J.GENE.2020.144422.

66 Dehaye, L., Duval, M., Viguier, D., Yaxley, J. and Job, D. (1997) Cloning and Expression of the Pea Gene Encoding SBP65, a Seed-Specific Biotinylated Protein. Plant Molecular Biology, Springer, 35, 605–621. 10.1023/A:1005836405211/METRICS.

67. Wilhelm, K.S. and Thomashow, M.F. (1993) Arabidopsis Thaliana Cor15b, an Apparent Homologue of Cor15a, Is Strongly Responsive to Cold and ABA, but Not Drought. Plant Molecular Biology, Kluwer Academic Publishers, 23, 1073–1077. 10.1007/BF00021822/METRICS.

68 Luo, D., Wu, Z., Bai, Q., Zhang, Y., Huang, M., Huang, Y. and Li, X. (2023) Universal Stress Proteins: From Gene to Function. International Journal of Molecular Sciences 2023, Vol. 24, Page 4725, Multidisciplinary Digital Publishing Institute, 24, 4725. 10.3390/IJMS24054725.

69 Kushwaha, R., Lloyd, T.D., Schäfermeyer, K.R., Kumar, S. and Downie, A.B. (2012) Identification of Late Embryogenesis Abundant (LEA) Protein Putative Interactors Using Phage Display. International Journal of Molecular Sciences 2012, Vol. 13, Pages 6582- 6603, Molecular Diversity Preservation International, 13, 6582–6603. 10.3390/IJMS13066582.

70 Grafahrend-Belau, E., Schreiber, F., Koschützki, D. and Junker, B.H. (2009) Flux Balance Analysis of Barley Seeds: A Computational Approach to Study Systemic Properties of Central Metabolism. Plant Physiology, Oxford Academic, 149, 585–598. 10.1104/PP.108.129635.

71 Shewry, P.R. and Halford, N.G. (2002) Cereal Seed Storage Proteins: Structures, Properties and Role in Grain Utilization. Journal of Experimental Botany, Oxford Academic, 53, 947– 958. 10.1093/JEXBOT/53.370.947.

72 Tanner, G.J., Colgrave, M.L., Blundell, M.J., Howitt, C.A. and Bacic, A. (2019) Hordein Accumulation in Developing Barley Grains. Frontiers in Plant Science, Frontiers Media S.A., 10, 443201. 10.3389/FPLS.2019.00649/BIBTEX.

73 Østergaard, O., Melchior, S., Roepstorff, P. and Svensson, B. (2002) Initial Proteome Analysis of Mature Barley Seeds and Malt. Proteomics, 2, 733–739. 10.1002/1615-9861(200206)2:6<733::aid-prot733>3.0.co;2-e.

74 Gorjanović, S. (2009) A Review: Biological and Technological Functions of Barley Seed Pathogenesis-Related Proteins (PRs). Journal of the Institute of Brewing, John Wiley & Sons, Ltd, 115, 334–360. 10.1002/J.2050-0416.2009.TB00389.X.

75 Graham, I.A. (2008) Seed Storage Oil Mobilization. Annual Review of Plant Biology, Annual Reviews, 59, 115–142. 10.1146/ANNUREV.ARPLANT.59.032607.092938/CITE/REFWORKS.

76. Vinje, M.A., Henson, C.A., Duke, S.H., Simmons, C.H., Le, K., Hall, E. and Hirsch, C.D. (2021) Description and Functional Analysis of the Transcriptome from Malting Barley. Genomics, Academic Press, 113, 3310–3324. 10.1016/J.YGENO.2021.07.011.

77 Schmitt, M.R., Skadsen, R.W. and Budde, A.D. (2013) Protein Mobilization and Malting- Specific Proteinase Expression during Barley Germination. Journal of Cereal Science, Academic Press, 58, 324–332. 10.1016/J.JCS.2013.05.007.

78 Viswanath, K.K., Varakumar, P., Pamuru, R.R., Basha, S.J., Mehta, S. and Rao, A.D. (2020) Plant Lipoxygenases and Their Role in Plant Physiology. Journal of Plant Biology, Springer, 63, 83–95. 10.1007/S12374-020-09241-X/FIGURES/3.

79 Hassan, M.J., Geng, W., Zeng, W., Raza, M.A., Khan, I., Iqbal, M.Z., Peng, Y., Zhu, Y. and Li, Z. (2021) Diethyl Aminoethyl Hexanoate Priming Ameliorates Seed Germination via Involvement in Hormonal Changes, Osmotic Adjustment, and Dehydrins Accumulation in White Clover Under Drought Stress. Frontiers in Plant Science, Frontiers Media S.A., 12, 709187. 10.3389/FPLS.2021.709187/BIBTEX.

80 Ma, Z., Marsolais, F., Bykova, N. V. and Igamberdiev, A.U. (2016) Nitric Oxide and Reactive Oxygen Species Mediate Metabolic Changes in Barley Seed Embryo during Germination. Frontiers in Plant Science, Frontiers Research Foundation, 7, 179637. 10.3389/FPLS.2016.00138/BIBTEX.

81 Bahmani, M., O’Lone, C.E., Juhász, A., Nye-Wood, M., Dunn, H., Edwards, I.B. and Colgrave, M.L. (2021) Application of Mass Spectrometry-Based Proteomics to Barley Research. Journal of Agricultural and Food Chemistry, American Chemical Society, 69, 8591–8609. 10.1021/ACS.JAFC.1C01871/ASSET/IMAGES/LARGE/JF1C01871_0002.JPEG.

82 Osama, S.K., Kerr, E.D., Yousif, A.M., Phung, T.K., Kelly, A.M., Fox, G.P. and Schulz, B.L. (2021) Proteomics Reveals Commitment to Germination in Barley Seeds Is Marked by Loss of Stress Response Proteins and Mobilisation of Nutrient Reservoirs. Journal of Proteomics, Elsevier, 242, 104221. 10.1016/J.JPROT.2021.104221.

83 Zaidi, I., Hanin, M., Saidi, M.N., Soltani, N. and Brini, F. (2024) The Barley Dehydrin 4 and Stress Tolerance: From Gene to Function. Environmental and Experimental Botany, Elsevier, 224, 105802. 10.1016/J.ENVEXPBOT.2024.105802.

84 Johnson, K.A., Kim, E., Teeter, M.M., Suh, S.W. and Stec, B. (2005) Crystal Structure of α- Hordothionin at 1.9 Å Resolution. FEBS Letters, John Wiley & Sons, Ltd, 579, 2301–2306. 10.1016/J.FEBSLET.2004.12.100.

85 Breiteneder, H. (2004) Thaumatin-like Proteins – a New Family of Pollen and Fruit Allergens. Allergy, John Wiley & Sons, Ltd, 59, 479–481. 10.1046/J.1398-9995.2003.00421.X.

86 Sampedro, J. and Cosgrove, D.J. (2005) The Expansin Superfamily. Genome Biology, BioMed Central, 6, 1–11. 10.1186/GB-2005-6-12-242/FIGURES/8.

87 Breiteneder, H. and Radauer, C. (2004) A Classification of Plant Food Allergens. Journal of Allergy and Clinical Immunology, Mosby Inc., 113, 821–830. 10.1016/J.JACI.2004.01.779/ASSET/F3E25BD8-CD50-4738-BF1C-2D5C27DB6695/MAIN.ASSETS/GR3.JPG.

88 Samtiya, M., Aluko, R.E. and Dhewa, T. (2020) Plant Food Anti-Nutritional Factors and Their Reduction Strategies: An Overview. Food Production, Processing and Nutrition 2020 2:1, BioMed Central, 2, 1–14. 10.1186/S43014-020-0020-5.

89 Deifl, S., Zwicker, C., Vejvar, E., Kitzmüller, C., Gadermaier, G., Nagl, B., Vrtala, S., Briza, P., Zlabinger, G.J., Jahn-Schmid, B., Ferreira, F. and Bohle, B. (2014) Glutathione-S- Transferase: A Minor Allergen in Birch Pollen Due to Limited Release from Hydrated Pollen. PLOS ONE, Public Library of Science, 9, e109075. 10.1371/JOURNAL.PONE.0109075.

90 Kerr, E.D., Caboche, C.H., Josh, P. and Schulz, B.L. (2021) Benchtop Micro-Mashing: High- Throughput, Robust, Experimental Beer Brewing. Scientific Reports 2021 11:1, Nature Publishing Group, 11, 1–10. 10.1038/s41598-020-80442-7.

91 Breiteneder, H. and Radauer, C. (2004) A Classification of Plant Food Allergens. Journal of Allergy and Clinical Immunology, Mosby, 113, 821–830. 10.1016/J.JACI.2004.01.779.

92 Bi, X., Ye, L., Lau, A., Kok, Y.J., Zheng, L., Ng, D., Tan, K., Ow, D., Ananta, E., Vafiadi, C. and Muller, J. (2018) Proteomic Profiling of Barley Spent Grains Guides Enzymatic Solubilization of the Remaining Proteins. Applied Microbiology and Biotechnology, Springer Verlag, 102, 4159–4170. 10.1007/S00253-018-8886-8/FIGURES/3.

93. Evans, D.E. and Bamforth, C.W. (2009) Beer Foam: Achieving a Suitable Head. Beer: A Quality Perspective, Academic Press, 1–60. 10.1016/B978-0-12-669201-3.00001-4.

94 Jaeger, A., Zannini, E., Sahin, A.W. and Arendt, E.K. (2021) Barley Protein Properties, Extraction and Applications, with a Focus on Brewers’ Spent Grain Protein. Foods 2021, Vol. 10, Page 1389, Multidisciplinary Digital Publishing Institute, 10, 1389. 10.3390/FOODS10061389.

95 Huang, Y., Jonsson, N.N., McLaughlin, M., Burchmore, R., Johnson, P.C.D., Jones, R.O., McGill, S., Brady, N., Weidt, S. and Eckersall, P.D. (2023) Quantitative TMT-Based Proteomics Revealing Host, Dietary and Microbial Proteins in Bovine Faeces Including Barley Serpin Z4, a Prominent Component in the Head of Beer. Journal of Proteomics, Elsevier, 285, 104941. 10.1016/J.JPROT.2023.104941.

96 Evans, D.E., Sheehan, M.C. and Stewart, D.C. (1999) The Impact of Malt Derived Proteins on Beer Foam Quality. Part II: The Influence of Malt Foam-Positive Proteins and Non-Starch Polysaccharides on Beer Foam Quality. Journal of the Institute of Brewing, John Wiley & Sons, Ltd, 105, 171–178. 10.1002/J.2050-0416.1999.TB00016.X.

97 Tatham, A.S. and Shewry, P.R. (2008) Allergens to Wheat and Related Cereals. Clinical & Experimental Allergy, John Wiley & Sons, Ltd, 38, 1712–1726. 10.1111/J.1365-2222.2008.03101.X.

98. Sander, I., Rozynek, P., Rihs, H.P., Van Kampen, V., Chew, F.T., Lee, W.S., Kotschy-Lang, N., Merget, R., Brüning, T. and Raulf-Heimsoth, M. (2011) Multiple Wheat Flour Allergens and Cross-Reactive Carbohydrate Determinants Bind IgE in Baker’s Asthma. Allergy, John Wiley & Sons, Ltd, 66, 1208–1215. 10.1111/J.1398-9995.2011.02636.X.

