## Supplementary information for "What’s left from the brew? Investigating residual barley proteins in spent grains for downstream valorization opportunities"

**for**

### Content of supplementary material

#### Supplementary tables

Table S1: Overview of raw data for dry matter, crude protein, and amino acid composition of barley, malt, and BSG.

Table S2: Appended .xlsx file covering MaxQuant output data and additional downstream processing hereof.

Table S3: BLAST analysis of differential proteins from pair-wise comparison of barley and malt.

Table S4: BLAST analysis of differential proteins from pair-wise comparison of malt and BSG.

Table S5: Predicted subcellular localization of all identified proteins using DeepLoc 2.0.

Table S6: Overview of the most abundant (riBAQ > 0.5%) proteins in any of the analyzed samples.

#### Supplementary figures

Figure S1: Scree Plot from principal component analysis of MaxLFQ data in MassDynamics.

Figure S2: Statistical analysis of subcellular protein localization.

Figure S3: Volcano-plot from pair-wise differential analysis of protein LFQ intensity for BSG vs. Cryo-BSG.

**Table S1:** Overview of raw data for dry matter, crude protein, and amino acid composition of barley, malt, and BSG. Amino acid composition is given as both mg/g dry matter basis and as relative (% w/w). Values are given as mean ± standard deviation based on three replications.

| Sample | Barley | Malt | BSG |
|---|---|---|---|
| **Dry Matter (%)** | 86.71 ± .04 | 94.53 ± .07 | 95.03 ± .03 |
| **Crude Protein (% DM)** | 8.99 ± .50 | 10.46 ± .80 | 17.42 ± .24 |
| | **Amino Acid Composition (mg/g DM)** | | |
| **PHE** | 2.89 ± .09 | 3.10 ± .06 | 7.05 ± .51 |
| **LEU** | 12.35 ± .42 | 13.16 ± .47 | 9.62 ± .16 |
| **ILE** | 3.25 ± .17 | 3.67 ± .07 | 5.47 ± .09 |
| **MET** | .00 ± .00 | .00 ± .00 | 1.31 ± .06 |
| **TYR** | 1.56 ± .23 | 1.96 ± .45 | 5.40 ± .42 |
| **PRO** | 8.70 ± .42 | 10.56 ± .25 | 16.62 ± .80 |
| **VAL** | 4.32 ± .07 | 4.90 ± .12 | 9.95 ± .70 |
| **ALA** | 4.01 ± .09 | 4.71 ± .16 | 8.10 ± .25 |
| **THR** | 2.67 ± .18 | 3.05 ± .04 | 5.73 ± .10 |
| **GLY** | 4.12 ± .69 | 3.93 ± .30 | 6.98 ± .27 |
| **SER** | 3.87 ± .24 | 3.95 ± .45 | 6.51 ± .20 |
| **HIS** | 2.82 ± .01 | 3.19 ± .10 | 7.12 ± .04 |
| **ARG** | 6.79 ± .30 | 7.48 ± .18 | 3.77 ± .09 |
| **LYS** | 3.27 ± .06 | 3.67 ± .35 | 5.60 ± 3.22 |
| **GLU/GLN** | 18.89 ± .86 | 18.31 ± .10 | 38.97 ± .68 |
| **C-C** | .00 ± .00 | .00 ± .00 | 5.73 ± .33 |
| **ASP/ASN** | 5.98 ± .23 | 6.04 ± .49 | 14.25 ± .84 |
| **ΣAA** | 85.49 ± .98 | 91.67 ± 1.11 | 158.18 ± 5.70 |
| **ΣEAA[1]** | 31.57 ± .21 | 34.73 ± .36 | 51.85 ± 4.56 |
| **ΣNEAA[2]** | 53.92 ± 1.00 | 56.95 ± .92 | 106.33 ± 1.14 |
| | **Amino Acid Composition (% (w/w))** | | |
| **PHE** | 3.39 ± .10 | 3.38 ± .03 | 4.45 ± .16 |
| **LEU** | 14.45 ± .59 | 14.36 ± .56 | 6.08 ± .11 |
| **ILE** | 3.80 ± .21 | 4.00 ± .13 | 3.46 ± .07 |
| **MET** | .00 ± .00 | .00 ± .00 | .83 ± .07 |
| **TYR** | 1.82 ± .28 | 2.13 ± .48 | 3.42 ± .13 |
| **PRO** | 10.17 ± .40 | 11.52 ± .22 | 10.51 ± .13 |
| **VAL** | 5.05 ± .04 | 5.34 ± .15 | 6.29 ± .21 |
| **ALA** | 4.69 ± .05 | 5.14 ± .11 | 5.12 ± .02 |
| **THR** | 3.12 ± .18 | 3.32 ± .04 | 3.63 ± .20 |
| **GLY** | 4.82 ± .86 | 4.29 ± .29 | 4.43 ± .33 |
| **SER** | 4.53 ± .24 | 4.31 ± .44 | 4.12 ± .02 |
| **HIS** | 3.30 ± .04 | 3.48 ± .06 | 4.50 ± .14 |
| **ARG** | 7.94 ± .28 | 8.16 ± .10 | 2.39 ± .14 |
| **LYS** | 3.83 ± .10 | 4.00 ± .35 | 3.51 ± 1.91 |
| **GLU/GLN** | 22.10 ± .86 | 19.98 ± .33 | 24.66 ± 1.32 |
| **C-C** | .00 ± .00 | .00 ± .00 | 3.63 ± .34 |
| **ASP/ASN** | 6.99 ± .20 | 6.59 ± .62 | 9.01 ± .21 |
| **ΣEAA[1]** | 36.93 ± .48 | 37.89 ± .35 | 32.75 ± 1.70 |
| **ΣNEAA[2]** | 63.07 ± .48 | 62.11 ± .35 | 67.26 ± 1.70 |

[1]The sum of essential amino acids (ΣEAA) represents Phenylalanine (Phe), Leucine (Leu), Isoleucine (Ile), Methionine (Met), Valine (Val), Threonine (Thr), Hisidine (His), and Lysine (Lys).

[2]The sum of non-essential amino acids (ΣEAA) represents Tyrosine (Tyr), Proline (Pro), Alanine (Ala), Glycine (Gly), Serine (Ser), Arginine (Arg), Glutamic acid/Glutamine (Glu/Gln), Cystine (C-C), and Aspartic acid/Asparagine (Asp/Asn).

**Table S2:** Appended .xlsx file covering MaxQuant output data and additional downstream processing hereof. The "proteinGroups" folder has been modified to include a range of additional data used in the manuscript covering calculation of relative iBAQ (riBAQ), replicate means, abundance and reproducibility filters, and predicted subcellular localization (based on Table S5). Common contaminants and false positives have been manually filtered. For a full data file with no modification, see the referenced project in the PRIDE data repository. In addition to the "proteinGroups" pane, the .xlsx file also contains BLAST results for pair-wise analysis of barley and malt (Table S3), BLAST results for pair-wise analysis of malt and BSG (Table S4), prediction of subcellular localization for all identified proteins (Table S5), as well as further analysis and definition of short names for the most abundant proteins across all samples (Table S6).

**Table S3:** BLAST analysis of differential proteins from pair-wise comparison of barley and malt. Proteins are listed with their ISBC_v2 AC#, p-value from pair-wise analysis of triplicate MaxLFQ intensities in MassDynamics, fold-change, adjusted p-value, Uniprot AC# of best BLAST ID, Identity (%) of the BLAST hit, associated e-value, and a note on protein function and potential literature DOIs. Proteins are sorted from highly enriched in malt (high negative fold change) to depleted in malt (high positive fold change). Table can be found in appended .xlsx file (Table S2) as "Table S3".

**Table S4:** BLAST analysis of differential proteins from pair-wise comparison of malt and BSG. Proteins are listed with their ISBC_v2 AC#, p-value from pair-wise analysis of triplicate MaxLFQ intensities in MassDynamics, fold-change, adjusted p-value, predicted subcellular localization by DeepLoc, Uniprot AC# of best BLAST ID, Identity (%) of the BLAST hit, associated e-value, and a note on protein function and potential literature DOIs. Proteins are sorted from highly enriched in BSG (high negative fold change) to depleted in BSG (high positive fold change). Table can be found in appended .xlsx file (Table S2) as "Table S4".

**Table S5:** Predicted subcellular localization of all identified proteins using DeepLoc 2.0. For all proteins, the table includes ISBC_v2 AC#, predicted localization (multiple allowed), identified signals, probability for localization within specific subcellular compartments, and the preferential localization (as determined by the highest single compartment probability). Table can be found in appended .xlsx file (Table S2) as "Table S5".

**Table S6:** Overview of the most abundant (riBAQ > 0.5%) proteins in any of the analyzed samples. The table includes the ISBC_v2 AC#, a long name (as per BLAST analysis), a short name (as used in Figure 5B), mean riBAQ abundance across all four sample types, log2 transform of mean riBAQ abundances, and a fold change comparison of BSG vs. Malt. Table can be found in appended .xlsx file (Table S2) as "Table S6".

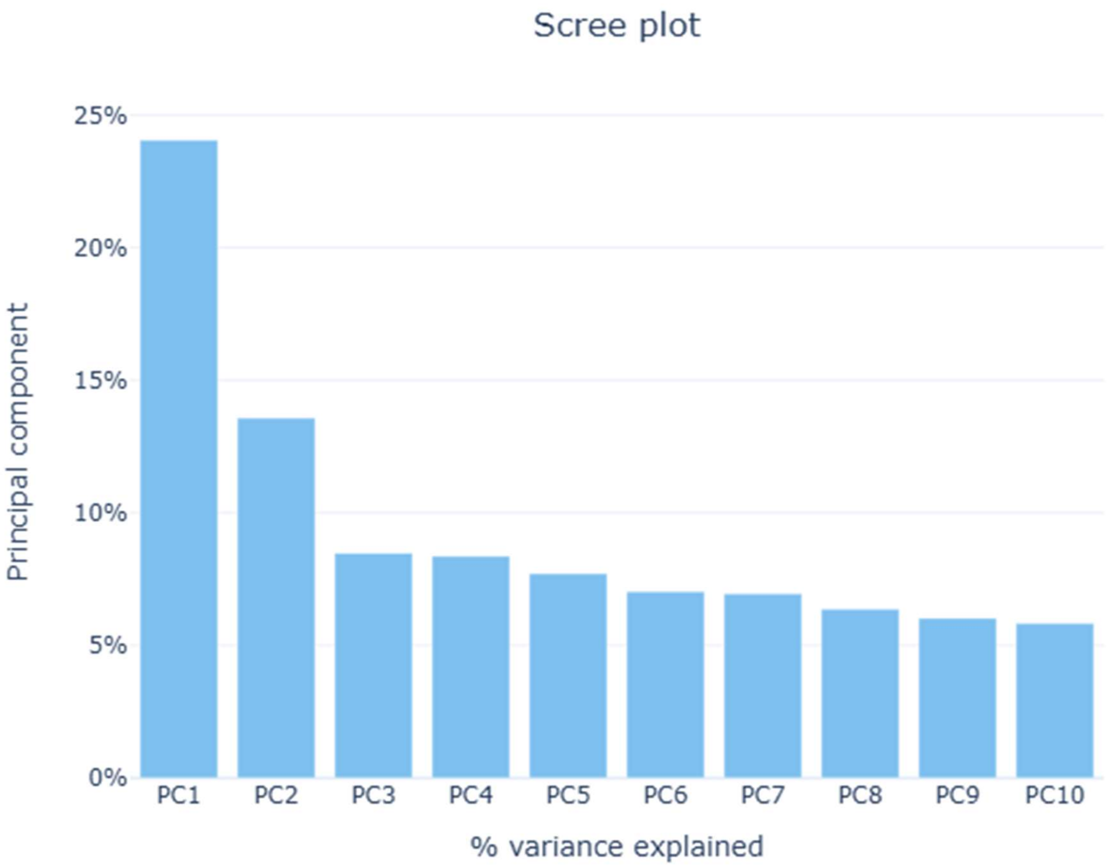


**Figure S1:** Scree Plot from principal component analysis of MaxLFQ data in MassDynamics. The scree plot shows the extent of variability explained by the first ten principal components, whereof the first two are used in Figure 3C.

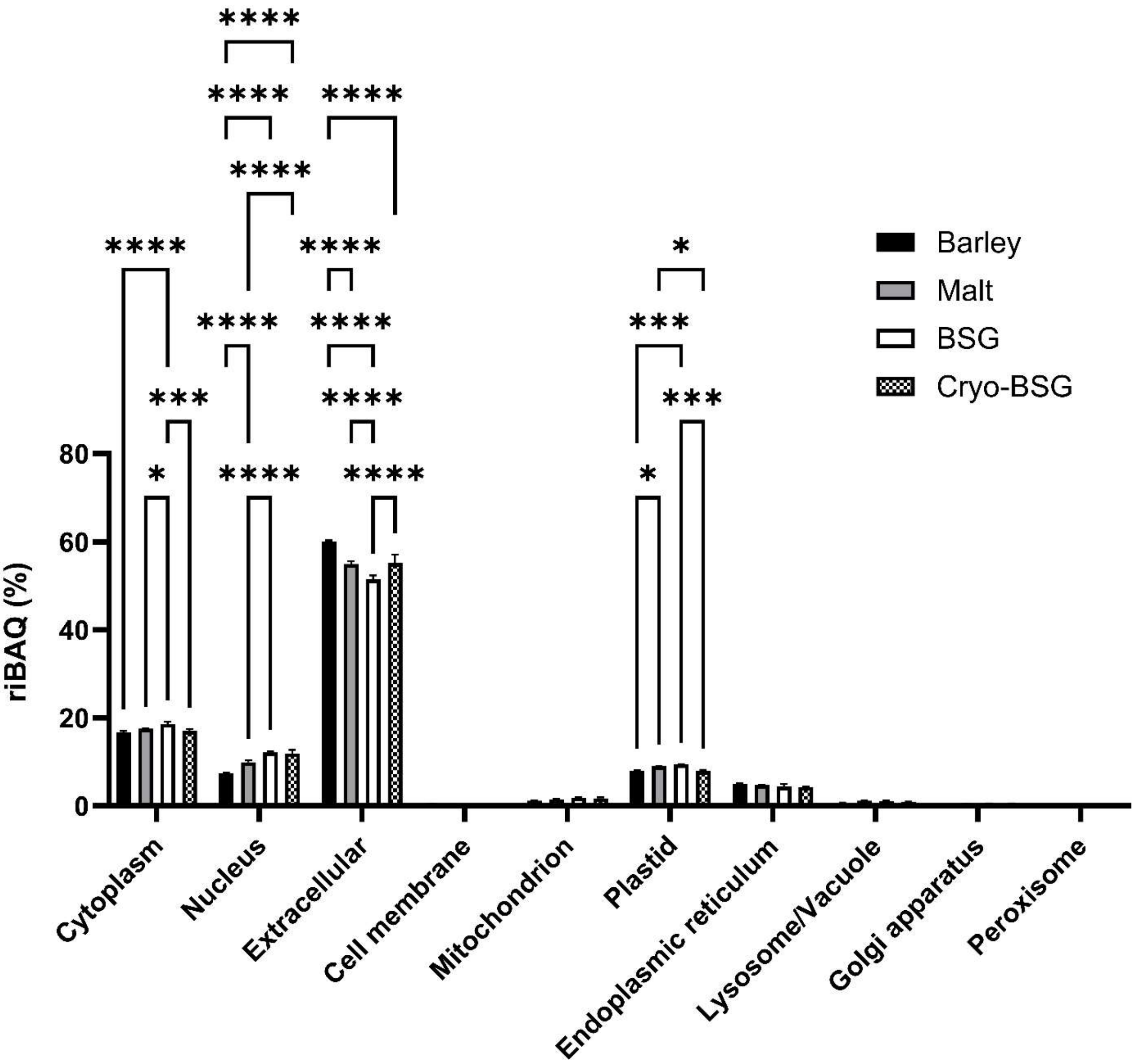


**Figure S2:** Statistical analysis of subcellular protein localization. The figure plots the same data as in Fig. 5A, but also includes statistical analysis of riBAQ distribution according to predicted subcellular localization using Tukey with a 95% confidence interval. Values are indicated as means with the standard deviation (n = 3). Statistical analysis is performed as two-way ANOVA with significance level (from adjusted p-values) indicated by "ns" (p > 0.05), "*" (p ≤ 0.05), "**" (p ≤ 0.01), "***" (p ≤ 0.001), and "****" (p ≤ 0.0001).

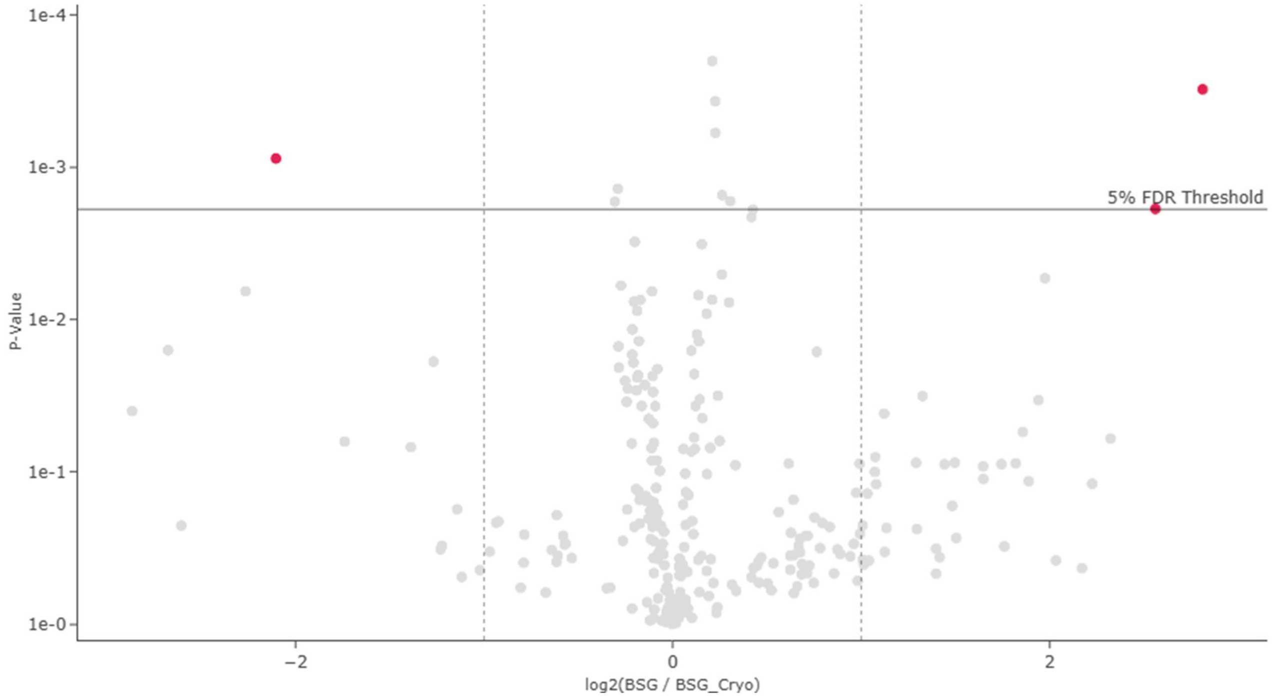


**Figure S3:** Volcano-plot from pair-wise differential analysis of protein LFQ intensity for BSG vs. Cryo-BSG. The analysis is performed in MassDynamics and highlighted proteins (red) were found to be significantly ($p < 0.05$) and substantially ($FC > 2$) differential in the pair-wise analysis.
